# Human internal exposures to alternariol and its monomethyl ether are predicted below thresholds of *in vitro* toxicity by physiologically based kinetic modeling

**DOI:** 10.64898/2026.05.11.724263

**Authors:** Eszter Borsos, Blandine Descamps, Nick Hetzschold, Elisabeth Varga, Doris Marko, Georg Aichinger

## Abstract

The foodborne mycotoxins alternariol (AOH) and alternariol monomethyl ether (AME) have been associated with several adverse effects, including cytotoxicity, genotoxicity, endocrine disruption, and immunomodulation. As these endpoints are typically observed *in vitro* at micromolar concentrations, the question arises whether such levels are attainable in exposed humans. To address this data gap in chemical risk assessment, a physiologically based kinetic (PBK) model was developed to predict internal exposure doses to AOH and AME in humans.

As input parameters, kinetic constants for hepatic glucuronidation were obtained *in vitro* by incubating Sprague Dawley rat and human liver S9 fractions with 0.5-50 µM AOH and 0.5-20 µM AME, demonstrating rapid biotransformation in both species. Intestinal absorption of AME and physicochemical parameters were estimated using quantitative structure-activity relationship (QSAR) models. Sensitivity analysis identified parameters describing hepatic glucuronidation and gastrointestinal uptake as among the most influential, confirming the importance of their reliable estimation. The PBK model was evaluated against available rodent toxicokinetic data and subsequently extrapolated to humans. Ultimately, the currently available exposure estimates published by EFSA in 2016 were applied to predict target tissue concentrations, which were compared to points of departure (PoDs) for relevant toxicological endpoints.

Even in the most susceptible group of male toddlers, predicted internal concentrations (10□□ µM range) were approximately four orders of magnitude below the respective PoDs. Consequently, under the applied exposure assumptions and considering the compounds as isolated chemicals, AOH and AME are not expected to reach systemic or tissue concentrations associated with the investigated effects.

## 1 Introduction

Dietary exposure to naturally occurring mycotoxins remains a persistent food safety concern worldwide, continuously posing new challenges through altered fungal distribution, crop susceptibility, and contamination patterns by climate change (Casu *et al*. 2024). Although regulatory efforts naturally focus on well-characterized and regulated mycotoxins, attention has increasingly shifted toward understudied compounds for which so far no regulation exists – so-called emerging mycotoxins. (Gruber-Dorninger *et al*. 2017).

Alternariol (AOH) and alternariol monomethyl ether (AME) fall into this category and are secondary metabolites of the ubiquitous fungal genus *Alternaria*, occurring in various food commodities, including grains, vegetables, fruits, legumes, oilseeds, nuts, and processed foodstuffs (EFSA 2011). These dibenzo-α-pyrones (structures shown in Figure 1) are relatively persistent, as substantial fractions can remain unchanged after heat treatments upon food processing and enzymatic digestion (Call *et al*. 2026; Estiarte *et al*. 2018).

**Figure 1:**
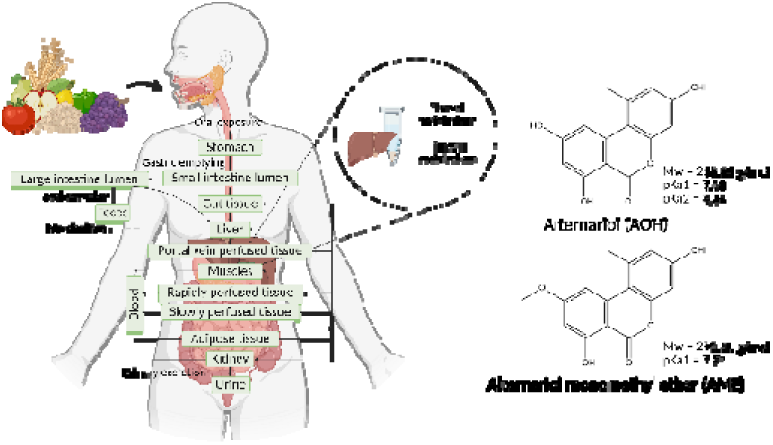
PBK model concept and chemical structures of AOH and AME. Created in BioRender. Aichinger, G. (2026) https://BioRender.com/aziuwd2.

Based on the available occurrence and toxicity data of AOH and AME, the European Food Safety Authority (EFSA) concluded that the estimated chronic dietary exposure may exceed the relevant threshold of toxicological concern in some population groups (EFSA 2016). Thus, the Commission Recommendation (EU) no. 2022/553 encourages monitoring the concentration of AOH and AME in food and set indicative levels in selected food matrices (European Commission 2022). However, no binding maximum levels for AOH and AME exist due to the lack of a comprehensive risk assessment. This regulatory gap can be explained by the limited available data (Aichinger *et al*. 2021; EFSA 2011; Louro *et al*. 2024).

Despite lacking legislation for AOH and AME, their broad occurrence paired with substantial structural stability warrants caution given the range of adverse effects reported in numerous *in vitro* and limited *in vivo* studies. In different mammalian cell systems, genotoxicity of both AOH and AME has been detected *via* DNA strand breaks and micronucleus formation (Lehmann *et al*. 2006; Pfeiffer *et al*. 2007; Solhaug *et al*. 2013; Tang *et al*. 2022). Mechanistically, AOH and – to a lesser extent – AME act as topoisomerase I and II poisons, thereby initiating DNA damage responses (Fehr *et al*. 2009). Additionally, both toxins induce oxidative stress and apoptosis, which can contribute to genotoxicity and cytotoxicity, respectively (Bensassi *et al*. 2011; Tiessen *et al*. 2013). Furthermore, AOH and presumably AME display endocrine-disrupting effects, acting as weak estrogen receptor agonists *in vitro* (Lehmann *et al*. 2006; Dellafiora *et al*. 2018). In intestinal and immune-relevant cell lines, also immunomodulatory and pro-inflammatory effects of AOH and AME have been observed (Solhaug *et al*. 2015; Grover and Lawrence 2017; Schmutz *et al*. 2019). Given their frequent co-occurrence, these adverse effects might be further exacerbated through additive or synergistic interactions between AOH, AME, and other *Alternaria* toxins (Hohenbichler *et al*. 2020; Marin *et al*. 2024).

Consequently, one might wonder whether realistic AOH or AME exposure could lead to these adverse effects in humans. For this purpose, physiologically based kinetic (PBK) modeling provides a valuable *in silico* tool to predict the absorption, distribution, metabolism, and excretion (ADME) of a chemical, by incorporating physiological and anatomical, as well as physicochemical and kinetic parameters into a set of differential equations (Rietjens *et al*. 2011). This method facilitates quantitative *in vitro* to *in vivo* extrapolation (QIVIVE), enabling prediction of internal concentrations without the need to generate new animal data. Thus, the application of such new approach methodologies (NAMs) for the risk assessment of xenobiotics aligns with the global aim of replacing, reducing, and refining (3Rs) animal experiments (Paini *et al*. 2019).

In this study, we aimed to develop a PBK model predicting time-resolved tissue concentrations of AOH and AME in humans after oral intake across realistic exposure scenarios and population groups. First, a rodent model for AOH was developed and evaluated. The human PBK framework was then developed *via* cross-species extrapolation and subsequently extended to AME by structure-specific parameterization. The required physiological, physicochemical, and kinetic parameters were obtained from literature or derived from quantitative structure-activity relationship (QSAR) predictions. Additionally, *in vitro* incubations were performed to acquire data on the uridine diphosphate glucuronosyltransferase (UGT)-mediated glucuronidation kinetics of both toxins. The respective PBK models were used to translate known exposure data (EFSA 2016) of different population groups into predicted unbound concentrations in blood and target tissues. We then compared these values with *in vitro* points-of-departure (PoDs) for relevant endpoints to evaluate whether tissue concentrations may approach or exceed those PoDs, thereby fusing the available data to provide a quantitative, integrated understanding of the bioactivity of AOH and AME as a preliminary step towards risk assessment.

## 2 Materials and Methods

### 2.1 Glucuronidation of AOH and AME in rat and human liver S9 fractions

#### Chemicals and biological materials

Liver S9 fractions prepared from a pool of male Sprague Dawley rats (n = 25), as well as pooled human liver S9 fractions from mixed-gender donors (n = 6) were purchased from Gibco™ (Thermo Fisher Scientific Inc., Waltham, MA, USA). β-glucuronidase from *E*. *coli* was obtained from Sigma-Aldrich (Merck GmbH, Darmstadt, Germany).

Dimethyl sulfoxide (DMSO) (Carl Roth GmbH & Co., Karlsruhe, Germany) was used as a solvent for the toxin stock solutions. AOH (purity min. 95%) was purchased from Biomol GmbH (Hamburg, Germany). AME was synthesized according to Mikula *et al*. (2013) and kindly provided by Prof. Roderich Süssmuth (Technical University of Berlin, Berlin, Germany). Further information regarding this synthesis route is provided in the supplementary information. To ensure reproducibility within the incubation studies, Gentest™ UDPGA (uridine diphosphate glucuronic acid; solution A) and uridine 5′-diphospho-glucuronosyltransferase (UGT) buffer (solution B) were purchased from a commercial source (Discovery Life Sciences, Woburn, MA, USA). Anhydrous sodium acetate (min. 99%) was obtained from Honeywell Specialty Chemicals (Seelze, Germany).

For the sample preparation and analysis with liquid chromatography coupled to mass spectrometry (LC-MS), CHROMASOLV LC–MS-grade acetonitrile and methanol (Honeywell Riedel-de Haën, Seelze, Germany) was used. HiPerSolv CHROMANORM LC–MS-grade water was purchased from VWR International (Radnor, PA, USA). As mobile phase additives, ammonium acetate and ammonium hydroxide (both LC-MS grade) were purchased from Sigma-Aldrich (St. Louis, MO, USA). Certified analytical standards of AOH and AME were obtained from Romer Labs Diagnostic GmbH (Tulln, Austria).

#### Incubation with rat and human liver S9 fractions to obtain kinetic information regarding glucuronidation

The glucuronidation kinetics of the target toxins were studied by incubating rat and human liver S9 fractions with 0.5, 1, 2, 5, 10, and 20 µM AOH or AME. The protocol by Ren *et al*. (2024) was adapted according to the manufacturer’s recommendations for the Gentest^TM^ UDPGA and UGT buffer.

Preliminary studies were performed to optimize incubation time and protein concentration. Due to the rapid glucuronidation observed in both species, a short incubation time of 3 min was selected. To maintain measurable levels of the residual parent toxin despite the rapid metabolic conversion, relatively low S9 protein concentrations were used. The final protein concentrations were 0.05 mg/mL (rat liver S9) and 0.2 mg/mL protein (human liver S9), as well as 0.35 mg/mL (rat liver S9) and 0.075 mg/mL protein (human liver S9) for AOH and AME, respectively.

The preincubation mixture contained the toxin of interest in DMSO and the S9 fraction dissolved in UGT buffer (solution B). After a 2-minute preincubation at 37 °C (mixing at 300 rpm), UDPGA was added to initiate the metabolic reaction. The final incubation mixture contained 1% (*v/v*) DMSO and 1× solution B, corresponding to 50 mM Tris-HCl (pH 7.5), 8 mM MgCl_2_, 25 µg/mL alamethicin, and 2 mM UDPGA.

After the incubation time of 3 min had passed, 50 µL of ice-cold extraction solvent (acetonitrile–methanol, 1:1, *v/v*) was added to the incubation solution to terminate the reaction. Subsequently, the samples were placed at −20 °C for at least 1 h to ensure protein precipitation, centrifuged (15 min, 18,000 × *g*, 4 °C), and the supernatant was further diluted with methanol–water (1:1, *v/v*) prior to LC–MS analysis. In the solvent control, the toxin solution was substituted with DMSO in the incubation mixture. Control incubations were performed by replacing the cofactor UDPGA (solution A) with deionized water.

As an additional control for glucuronide formation, 50 µL of the non-terminated incubation mixture was immediately pipetted into 50 µL of β-glucuronidase solution (0.1 M sodium acetate buffer, pH 5, 1000 units/mL) and incubated for 1 h at 37 °C with shaking at 300 rpm. This enzymatic treatment hydrolyzes glucuronide conjugates to the corresponding parent compound, thereby confirming the identity of the detected metabolites as glucuronides. Subsequently, the reaction was terminated by adding 100 µL of ice-cold extraction solvent (acetonitrile–methanol, 1:1, *v/v*), and the same sample preparation procedure was performed as described above. The subsequent LC-MS/MS analysis was performed based on a published procedure by Borsos *et al*. (2024). Further details are provided in Supplementary Information 1.

#### Determination of kinetic constants as input parameters in the PBK model

As the parent compounds could only be reliably quantified in the S9 fraction incubation samples, kinetic calculations were based on substrate depletion rather than metabolite formation rates. The acquired concentration values were tested for normality, and outliers were detected and excluded according to the Nalimov test. Based on the quantified AOH or AME decrease, and considering the incubation time of 3 min, as well as the respective S9 fraction protein contents, transformation rates were calculated, which were then depicted against the treatment concentration for curve fitting. A standard Michaelis-Menten equation was fitted to these data to obtain the kinetic parameters (1). To this end, the integrated toolbox of OriginPro (v. 10.2.0.188., OriginLab Corporation, Northampton, MA, USA) was applied with orthogonal distance regression without weighing of the x values.

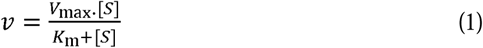

where [S] is the substrate concentration in µM, v is the rate of substrate depletion (nmol/min/mg S9 protein), V_max_ is the apparent maximum transformation rate (nmol/min/mg S9 protein), and K_m_ represents the Michaelis-Menten constant in µM.

### 2.2 PBK modeling

To address the data gap regarding the toxicokinetics of AOH and AME, we developed a rodent PBK model and evaluated its predictive accuracy against previously published animal toxicokinetic data (Schuchardt *et al*. 2014). Subsequently, the model was extrapolated to human physiology and finally extended to predict AME kinetics within humans using QSAR methods for chemical-specific parameterization. The conceptual model structure was adapted from a previously published human PBK model for urolithin A, a structurally related dietary chemical (Aichinger *et al*. 2023). As diet is the primary route of human exposure to these mycotoxins, the model was specifically designed for oral administration.

#### Model structure

The model structure describes oral uptake *via* the stomach lumen, followed by gastric emptying into the small and large intestine lumens, and subsequent fecal excretion. Absorption was modeled from the gut lumen into the gut tissue, followed by transport to the liver. Hepatic biotransformation was modeled to include phase I metabolism and phase II glucuronidation for both AOH and AME. Systemic circulation distributed the parent compounds to other organ tissues, with urinary excretion modeled *via* the kidneys by using the glomerular filtration rate (GFR) as well as the fraction unbound in plasma (FU) as factors describing the clearance velocity. When not individually compartmentalized, organs were combined into slowly perfused tissues (skin, bone), rapidly perfused tissues (heart, lung), or portal-vein-perfused tissues (spleen, pancreas, stomach). A specific uterus compartment was included for the human female models. All code was compiled and solved using Berkeley Madonna (version 10).

#### Model parameterization

Population-specific physiological parameters were collected from the literature (Table S2). Rodent parameters, including organ volume fractions and blood flows, were obtained from Brown *et al*. (1997) and from Crispens (1975). For the murine model, small (SI) and large intestine (LI) lumen surface areas as well as volume were calculated by assimilating the organs as cylinders (Casteleyn *et al*. 2010). Glomerular filtration rate value was obtained from Bivona *et al*. (2011). Murine gastric emptying rates and intestinal transit times were respectively derived from Schwarz *et al*. (2002) and Woting and Blaut (2018).

For human models, physiological parameters were predominantly sourced from the International Commission on Radiological Protection (Valentin 2002). Fractional organ volumes were computed, and the volumes of combined compartments were calculated as the sum of their constituent organ volumes. The surface areas of the small and large intestine lumens were calculated as described by Helander and Fändriks (2014), according to Equation (2)(1). Volumes of the small and large intestine lumens were calculated by assimilating the organs as cylinders (Valentin 2002).

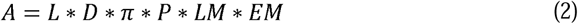

where L is the length of the intestine; D is the diameter of the intestine; P is the surface amplification due to plicae circulares; LM is the surface amplification due to villi observed by light microscopy; and EM is the surface amplification due to microvilli on the luminal surface of the absorptive cells observed by electron microscopy.

Glomerular filtration rate value was obtained from Levey *et al*. (2014). Physicochemical parameters included partition coefficients and intestinal uptake. *In silico* predictions of LogP and pK_a_ values (Table S3) for AOH and AME were generated using Chemaxon Playground (v1.6.2). Partition coefficients were calculated using the qivivetools toolbox (Punt *et al*. 2021), applying the algorithms of Berezhkovskiy (2004) for blood/tissue partitioning and Lobell and Sivarajah (2003) for plasma protein binding (Table S4). For the combined compartments, partition coefficients were computed with Equation (3).

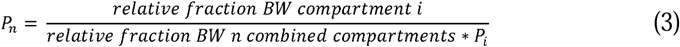

where P_n_ is the partition coefficient of n combined compartments, P_i_ is the partition coefficient of compartment i.

Due to the absence of experimental Caco-2 apparent permeability (P_app_) value for AME, this parameter was estimated using a relative scaling approach referenced to AOH. First, P_app_ values were predicted for both AOH and AME using two QSAR models (Kamiya *et al*. 2020; Lanevskij and Didziapetris 2019). These models yielded consistent predictions for each compound. A scaling factor was then calculated as the ratio of the available experimental *in vitro* AOH value (Burkhardt *et al*. 2009) to its mean QSAR-predicted value. This factor was applied to the mean QSAR prediction for AME to extrapolate its final calibrated P_app_ (Table S3). This coefficient is subsequently converted to *in vivo* effective permeability (P_eff_) using the regression for passively absorbed drugs by Sun *et al*. (2002).

Hepatic biotransformation phase I metabolism parameters for rodent and human were sourced from Borsos *et al*. (2024). Rates of phase I and phase II metabolism were scaled with microsomal and S9 scaling factors: rodent and human hepatic microsomal fractions with 62 mg/g liver (Smith *et al*. 2008) and 32 mg/g liver (Barter *et al*. 2007), respectively; rodent and human hepatic S9 fractions were scaled with 143 mg/g liver (Wang *et al*. 2020a) and 107.3 mg/g liver (Wang *et al*. 2020b), respectively.

#### Model evaluation and cross-species extrapolation

The rodent model (Table 1, Scenario 1) was designed to replicate the single oral administration of AOH to mice reported *in vivo* (Schuchardt *et al*. 2014). Model performance was evaluated by comparing predicted blood C_max_, t_max_, and Area Under the Curve (AUC) against the corresponding *in vivo* experimental data. Following evaluation, the rodent model was extrapolated to four human demographic groups (Scenarios 2–5) to represent the general population: adult male (73 kg body weight (bw)), adult female (60 kg bw), male child (19 kg bw), and male toddler (10 kg bw). As recently demonstrated by Bigonne *et al*. (2025) for Bisphenol A and several analogs, the comparison of specific physiological groups (sex, age) supports to reduce the traditional focus on male-only physiology, ensuring a more inclusive risk evaluation. Scenarios 6–9 applied the AME chemical parameters to these human physiological structures. Realistic human exposure doses (mean and 95th percentile, including lower and upper bounds) were derived from the latest ESFA report on *Alternaria* toxin exposure (EFSA 2016) to predict tissue-specific concentrations of toxicological interest.

**Table 1:**
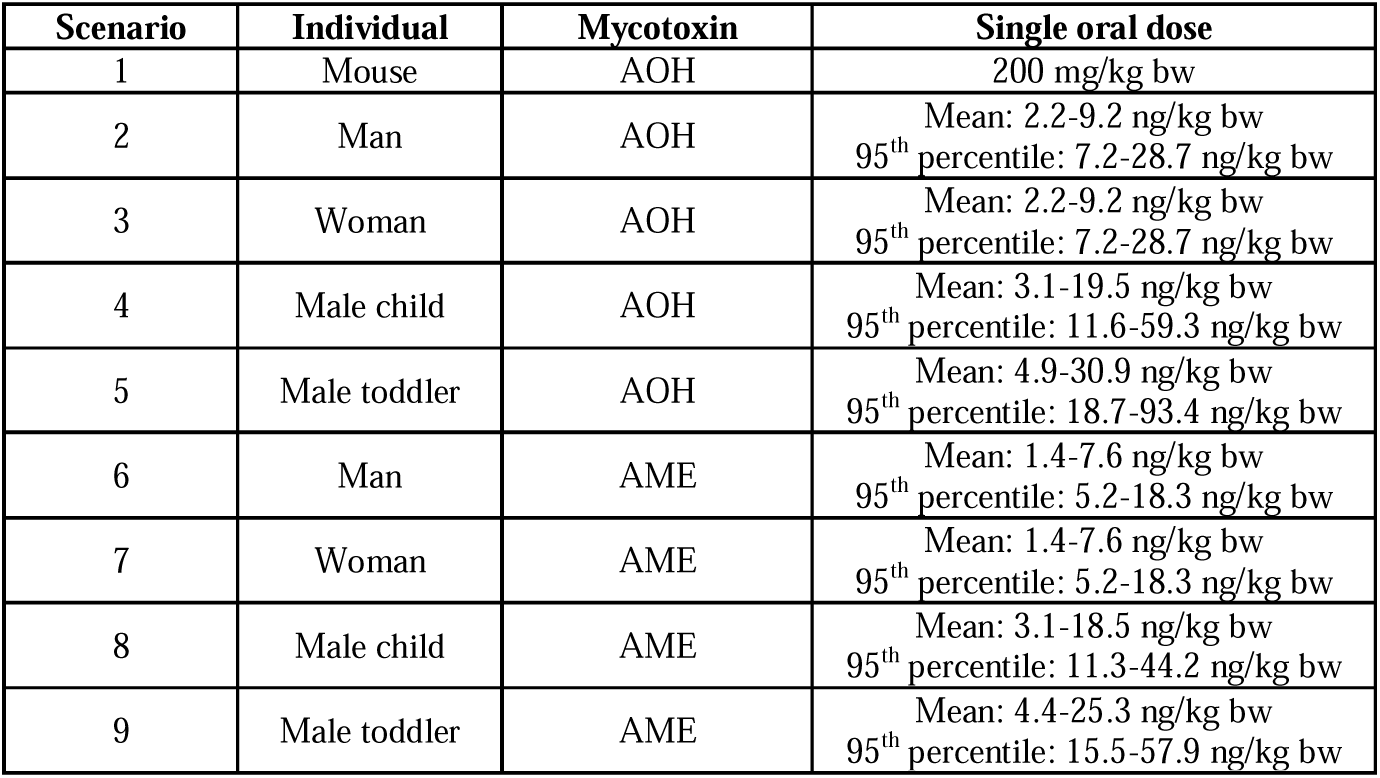
Exposure scenarios.

#### Sensitivity analysis

Local sensitivity analysis was performed to assess the influence of parameter deviations on model predictions, following the method of Evans (2000). Parameters were individually increased by 5%, with starting doses matching the human oral exposure scenarios. Normalized sensitivity coefficients (SC) were calculated using Equation (4). Parameters with an absolute |SC| > 0.1 were considered sensitive.

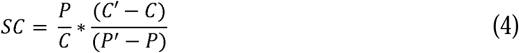

where C and C’ are the peak concentrations with raw and increased parameter, respectively, and P and P’ are the parameter values.

#### Monte-Carlo simulations

To account for inter-individual variability and parameter uncertainty, Monte Carlo (MC) simulations (n=500) were conducted focusing on the maximal concentration of AOH and AME in the blood for toddlers. Sensitive parameters identified in the previous step were sampled from probability distributions derived from the literature (Table S5). Coefficients of variation (CV) were calculated as ratio of standard deviation (σ) and mean (μ) for standard measurements or given usual value from literature. The integration interval was adjusted to 2.0*10^-4^ to ensure computational stability in Berkeley Madonna. Simulation outcomes were analyzed by comparing the median, 1^st^, and 3^rd^ quartiles of the concentration-time profiles.

#### Quantitative *in vitro* to *in vivo* extrapolation

We followed a forward dosimetry Quantitative *In Vitro* to *In Vivo* Extrapolation (QIVIVE) approach to characterize the potential health risks relative to single exposure of AOH and AME. Model-predicted peak concentrations (C_max_) in target organs (liver, kidney, colon, mammary tissues and uterus) were directly compared to Lowest Observed Adverse Effect Levels (LOAELs) obtained from the review by Louro *et al*. (2024). For each target tissue and single exposure time (24 h, 48 h, 72 h), we applied a “worst-case scenario” strategy. When multiple cell lines for the same endpoint (e.g., genotoxicity, immunotoxicity, or endocrine disruption) were reported for a single tissue, the lowest available LOAEL was selected as the threshold. The models were run to match the acute exposure time of the *in vitro* studies (24 h, 48 h, and 72 h). The Margin of Safety (MoS) was then calculated by calculating the ratio between the LOAEL and the predicted C_max_ of the respective target organ.

## 3 Results and discussion

### 3.1 *In vitro* glucuronidation kinetics

#### *In vitro* kinetic parameters

Incubations of rat and human liver S9 with AOH and AME in the presence of the cofactor UDPGA revealed a rapid and concentration-dependent decrease for both compounds across the examined species that followed Michaelis-Menten kinetics (Figure 2). As no significant depletion was observed in control incubations lacking UDPGA (Figures S1-S2), this is clearly attributable to UGT-dependent glucuronidation. Furthermore, two baseline-separated glucuronide peaks were observed for AOH and AME in their LC–MS/MS chromatograms (Figures S3 and S4, respectively), at the expected glucuronide MRM transitions. The corresponding signals at the parent MRM transition at the same retention times – particularly in the case of AME – point to partial in-source fragmentation of the glucuronides. These peaks appeared only in UDPGA-fortified S9 incubations and were abolished by post-incubation β-glucuronidase treatment, confirming the identities of the detected conjugates and supporting glucuronidation as the primary driver of the parent compound decrease under the applied assay conditions.

**Figure 2:**
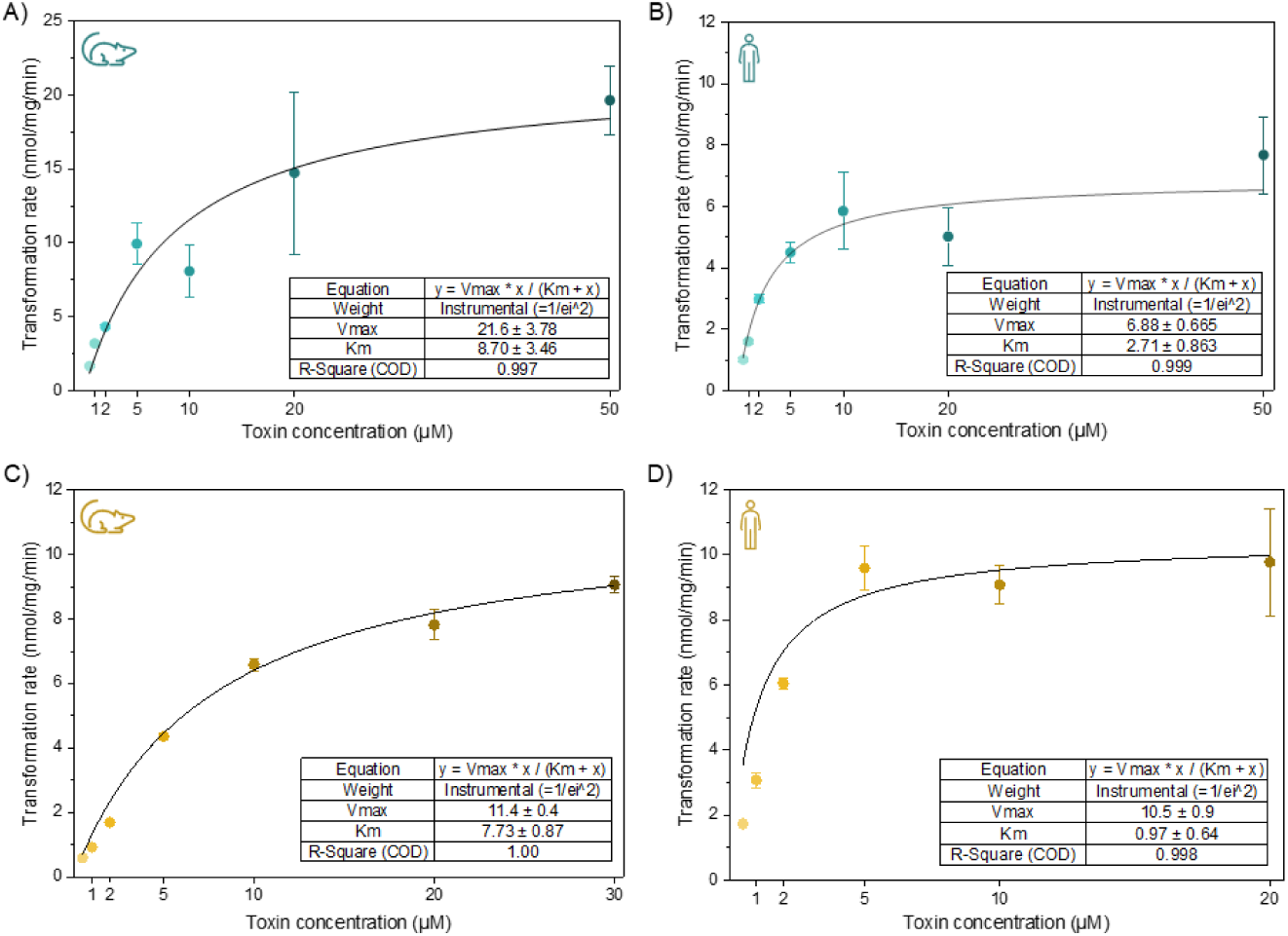
Concentration-dependent depletion of AOH. (A-B) and AME (C-D) in rat (A and C) and human (B and D) liver S9 fractions through glucuronidation in the presence of the cofactor UDPGA after an incubation time of 3 min. Data points represent means ± SD of four technical replicates. Michalis-Menten curves were fitted and respective parameters calculated with OriginPro by orthogonal distance regression without weighing of the x values. Please note that (A) has a different y-axis scaling than the other graphs.

For AOH, both unscaled kinetic parameters, *V*_max_ and *K*_m_, were higher in rat liver S9 fractions (**Figure 2**A) than human liver S9 (**Figure 2**B). Thus, human hepatic enzymes show a higher apparent affinity but lower capacity towards AOH compared to rats. Overall, the *in vitro* intrinsic clearance (*CL*_int_) values – calculated as the ratio of these kinetic parameters – yield approx. 2.5 mL/min/mg protein) in both rat and human S9 fractions (**Table 2**). This result suggests a comparable phase II metabolic stability of AOH across the species studied. However, after scaling these values to the *in vivo* situation using speciesLJspecific physiological scalars, the rat intrinsic clearance value (19.49 L/min/kg body weight; **Table 2**) is significantly higher than that in humans (8.12–10.35 L/min/kg body weight), indicating an approximately twofold more effective glucuronidation of AOH in rats *in vivo*. This pattern aligns with well-documented interspecies trends: rodents possess greater hepatic enzyme capacity per unit body weight than humans, yielding higher metabolic throughput (Martignoni *et al*. 2006). Nevertheless, because intrinsic clearance values exceeding 45 mL/min/kg (0.045 L/min/kg) are typically classified as high in humans (Vamsi Krishna *et al*. 2021), AOH can be still clearly defined as a high-clearance toxin in humans.

**Table 2:** Kinetic parameters for the glucuronidation of AOH and AME in incubations with UDPGA-fortified rat and human population scaled S9 fractions.

| Molecule name | Population/ S9 fractions | $V_{max}$ (nmol/mg/min) | $K_M$ (μM) | $CL_{int}$ (mL/min/mg) | VLS9 Scaling factor (mg S9 protein/g liver) | Scaled $CL_{int}$ (L/min/kgbw) |
| --- | --- | --- | --- | --- | --- | --- |
| Alternariol (AOH) | Rat | 21.6 | 8.70 | 2.48 | 143 | 19.49 |
|  | Human |  |  |  |  |  |
|  | Male | 6.88 | 2.71 | 2.54 | 107.3 | 8.69 |
|  | Female |  |  | 2.54 |  | 8.12 |
|  | Male child |  |  | 2.54 |  | 10.19 |
|  | Male toddler |  |  | 2.54 |  | 10.35 |
| Alternariol monomethyl ether (AME) | Rat | 11.4 | 7.73 | 1.47 | 143 | 11.58 |
|  | Human |  |  |  |  |  |
|  | Male | 10.5 | 0.97 | 10.8 | 107.3 | 37.05 |
|  | Female |  |  | 10.8 |  | 34.61 |
|  | Male child |  |  | 10.8 |  | 43.44 |
|  | Male toddler |  |  | 10.8 |  | 44.14 |

Due to the higher lipophilicity of AME (Figure 1, Table S4), only a concentration range of 0.5-20 µM was initially tested for this compound. However, because enzyme saturation was not achieved at 20 µM AME in rat hepatic S9 fractions, an additional substrate concentration of 30 µM was incorporated into the dataset to enhance the reliability of Michaelis-Menten curve fitting. The enzymatic capacities of rat and human liver UGTs were similar toward AME, with *V*_max_ values of 11.4 and 10.5 nmol/mg/min, respectively. However, substantial differences in the Michaelis constant were observed: *K*_m_ was 0.97 µM in human liver S9 fractions, whereas a higher value of 7.73 µM was measured in rat liver S9 fractions, implying a lower apparent affinity of rat hepatic enzymes for AME. Consequently, the calculated *in vitro* intrinsic clearance values also differ, pointing toward a more efficient glucuronidation of AME by humans than rats. These interspecies differences persist even after scaling the *in vitro* clearance values to *in vivo*: humans exhibit approximately fourfold more efficient glucuronidation of AME than rats (34.61–44.14 L/min/kg body weight vs 11.58 L/min/kg), classifying AME as a very highLJclearance compound in humans. A possible explanation is species differences in the UGT isoform profiles and expression between rats and humans and the fact that AOH and AME are metabolized by different UGT isoforms (Pfeiffer *et al*. 2009b). In humans, the isoform UGT2B15 show a higher contribution to the glucuronidation of AME compared to that of AOH, and the human liver expresses this isoform at substantial levels (Nakamura *et al*. 2008). In contrast, rat UGT2B isoforms are phylogenetically distinct from human UGT2B15, so that the lack of a direct functional equivalent might reflect less effective AME glucuronidation in rats (Nishihara 2014). Despite being a straightforward mechanistic hypothesis, because the specific rat UGT isoforms responsible for AOH and AME glucuronidation remain uncharacterized and metabolic enzyme functions cannot be directly translated between species, it cannot be directly tested with current data. Overall, the distinct speciesLJspecific trends between AOH and AME underscore the need to investigate structurally similar compounds individually, as their metabolic fate can differ fundamentally.

The assay conditions were selected to mitigate inherent limitations of the applied *in vitro* incubation approach. To enhance aqueous solubility of AOH and AME while minimizing solvent interference, the concentration of DMSO was kept at 1% (*v/v*). In a recent study, DMSO showed the least impact on the glucuronidation of 17β-estradiol among commonly used organic solvents (Wu *et al*. 2024). While solvent effects are substrate-, site-, and isozyme-specific, no publications have investigated these effects in the metabolism of our target compounds. Thus, estradiol – which shares UGT-relevant phenolic features to AOH and AME – served as a surrogate, providing initial guidance for solvent selection.

Both glucuronidation and sulfation have been reported as possible phase II conjugation pathways of AOH and AME in a range of *in vitro* systems (Pfeiffer *et al*. 2009a; Burkhardt *et al*. 2009, 2011). However, most assays provide only one cofactor and rely on systems with non-physiological UGT and sulfotransferase (SULT) expression, leaving the *in vivo* contribution of these biotransformation pathways insufficiently resolved. In general, UGTs exhibit lower substrate affinity but higher catalytic capacity compared to SULTs, which typically display higher affinity but lower capacity (Lai *et al*. 2022). For AOH and AME, the available evidence indicates that glucuronidation is the prevailing pathway across relevant exposure ranges: an *in vivo* study in piglets reported glucuronide conjugates of both toxins as predominant metabolites (den Hollander *et al*. 2025). While quantitative interspecies differences between pigs and humans are expected, the qualitative preference for glucuronidation is likely conserved. On this basis, the data collection in the present work focused on characterizing the kinetics of the glucuronidation of AOH and AME. Furthermore, because PBK modeling of the structurally related polyphenol urolithin A indicated that hepatic metabolism predominates over intestinal metabolism in humans (Aichinger *et al*. 2023), we prioritized hepatic glucuronidation kinetics as the principal data input.

### 3.2 Extrapolation of AME intestinal permeability

In the absence of experimental Caco-2 apparent permeability (P_app_) data for AME, a relative scaling approach was developed using AOH as a reference. This strategy relied on the structural similarity between the two mycotoxins, assuming that model-specific biases in the QSAR predictions would be proportional for both analogs.

Initial P_app_ predictions generated by the ‘QSAR K’ (Kamiya *et al*. 2020) and ‘QSAR L’ (Lanevskij and Didziapetris 2019) models showed high inter-model consistency for both compounds (Figure 3). By utilizing the ratio between the experimental *in vitro* AOH value and its mean QSAR-predicted value, a calibration factor was derived and applied to the AME predictions. The resulting calibrated log(P_app_) value of –5.11 for AME was then implemented in the PBK model.

**Figure 3:**
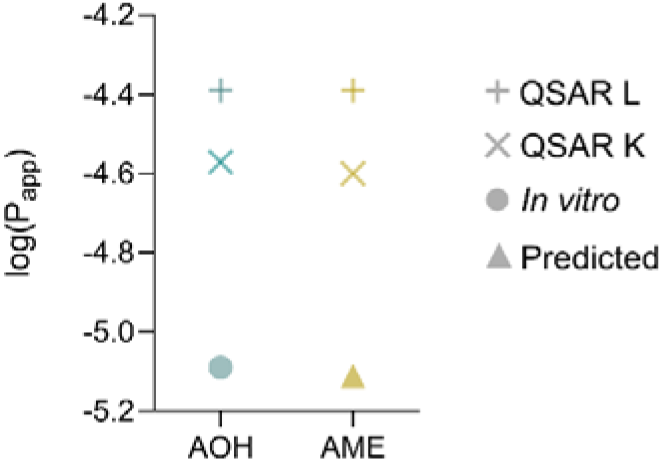
Prediction and relative scaling approach of Papp for AOH (blue) and AME (yellow) using a comparative QSAR approach. Initial Papp predictions values generated by the ‘QSAR K’ and ‘QSAR L’ models are depicted by plus and cross symbols respectively. *In vitro* value for AOH is depicted by the circle. Derivation of the final scaled Papp for AME is depicted by the triangle.

### 3.3 Rodent PBK model evaluation by comparison to *in vivo* data

Integrating the *in vitro* measured glucuronidation rates and scaled P_app_ to the parameterization, we developed a PBK model including passive diffusion across intestinal barriers, hepatic phase I and II (glucuronidation) metabolism, systemic tissue distribution, and renal clearance. Initially, the *in vivo* study from Puntscher *et al*. (2019), in which a complex *Alternaria* culture extract was administered to rats, was selected for evaluation. For this purpose, rat S9-derived glucuronidation rates were measured *in vitro*. However, the complex nature of the extract introduced significant mechanistic uncertainty, as competitive inhibition or synergistic potential effects among co-occurring toxins. We therefore parameterized a murine PBK model which was further evaluated against single-compound *in vivo* data reported by Schuchardt *et al*. (2014).

The use of rat-derived metabolic constants (V_max_, K_m_) within a mouse physiological framework represents an approximation justified by the absence of mouse-specific phase I and II metabolism data for AOH and AME. While both species share the major UGT enzyme families involved in xenobiotic metabolism, quantitative differences in enzyme activity and kinetic parameters between rats and mice have been documented for certain substrates (Shiratani *et al*. 2008). Nevertheless, rodents share physiological features that differ from those of humans, including a higher liver-to-body weight ratio (∼4–5% vs ∼2.5% in humans), greater intrinsic microsomal clearance, and a higher hepatic blood flow per kg of body weight (Kinatukara *et al*. 2026), supporting the hypothesis that inter-rodent metabolic differences are typically smaller than rodent-human differences. Thus, given that the primary objective was to validate the fundamental PBK structure and kinetic principles before human extrapolation, this approach provided a reasonable first approximation.

To evaluate the accuracy of the murine model, predicted plasma profiles were compared against *in vivo* mouse data following a single oral dose of 200 mg/kg bw AOH (Schuchardt *et al*. 2014). The predicted C_max_ and AUC values closely matched experimental results, while the predicted peak time was slightly accelerated (Figure 4, Table 3). Given that AOH is glucuronidated and structurally similar to urolithin A, which is known to undergo biliary excretion (Aichinger *et al*. 2023; Espín *et al*. 2007), we hypothesize that this temporal shift results from enterohepatic circulation (EHC) mediated by hepatic efflux transporters. AOH likely undergoes transporter-mediated hepatic efflux, followed by intestinal reabsorption, which is currently not implementable in the model due to a lack of respective QSAR tools to predict biliary excretion. This potentially causes a slight underestimation of systemic AOH concentrations with repeated exposure. The complete model code is available in Suppplementary informations S2-S10.

**Figure 4:**
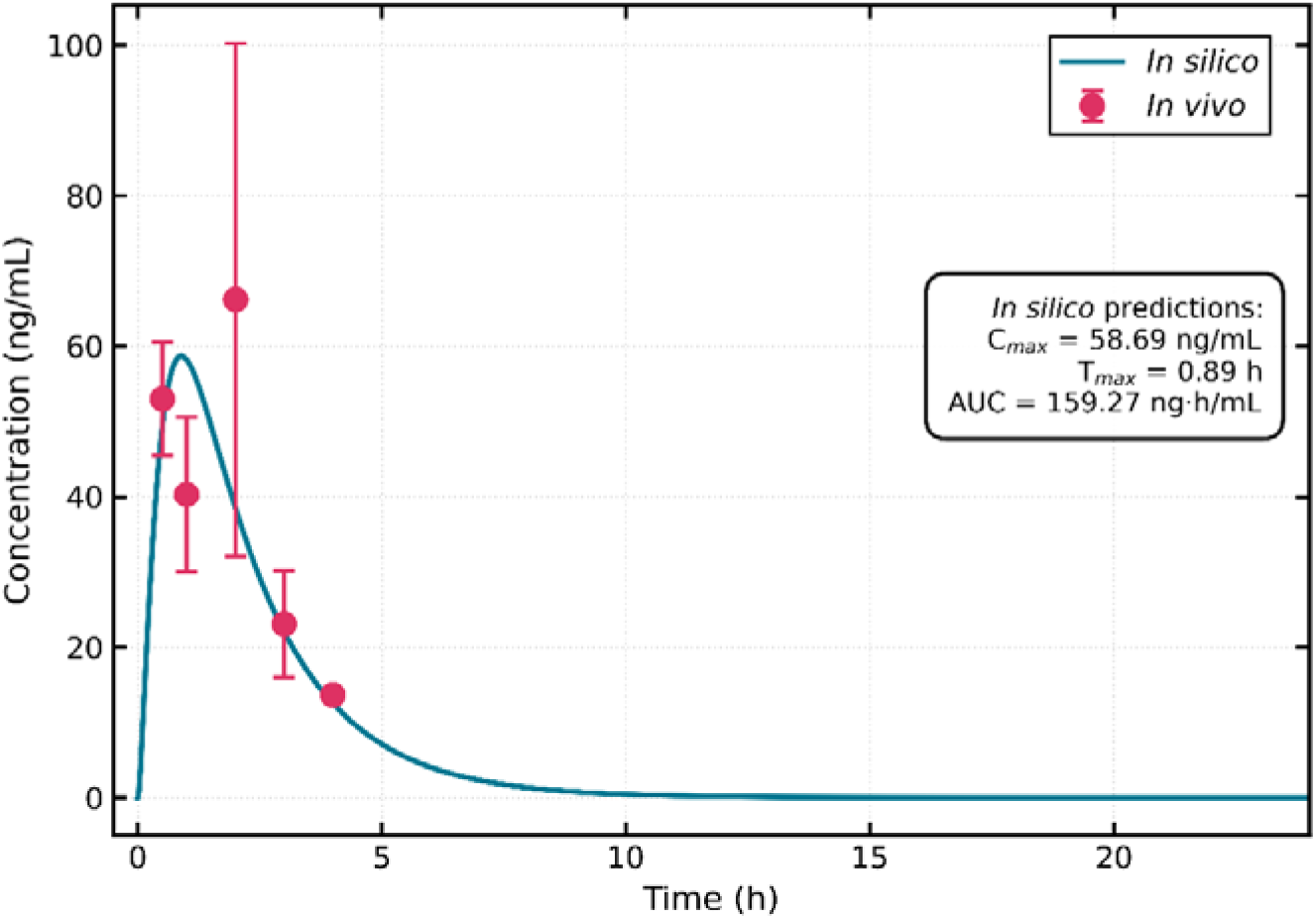
Predicted compared to measured blood concentrations of AOH after supplementation with 200 mg/kg bw AOH in mice. Red circles represent mean blood concentrations reported in the *in vivo* study by Schuchardt *et al*., 2014. Blue solid lines depict model predictions.

**Table 3:**
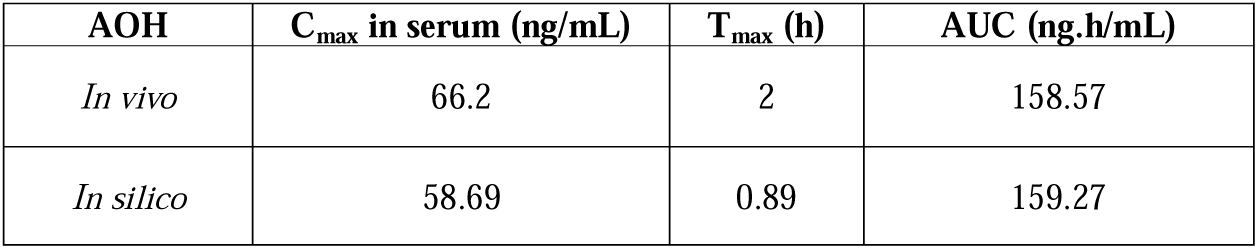
Rodent model evaluation for alternariol (AOH). AUC – area under the curve.

| AOH | C<sub>max</sub> in serum (ng/mL) | T<sub>max</sub> (h) | AUC (ng.h/mL) |
| --- | --- | --- | --- |
| *In vivo* | 66.2 | 2 | 158.57 |
| *In silico* | 58.69 | 0.89 | 159.27 |

### 3.4 Extrapolation to human models and internal exposure assessment

Following the evaluation, the murine model served as the basis for interspecies extrapolation to human physiology and metabolism. We subsequently extended this framework to include predictions of AME toxicokinetics. Four human demographic models, adult male, adult female, male child, and male toddler, were parameterized for each toxin. Concentrations of AOH and AME were predicted in the blood for all four demographic groups and exposure scenarios (Figure 5, Table 4). A consistent trend was observed across all exposure scenarios. On a body-weight-normalized basis, the internal exposure in toddlers was significantly higher than in adults, identifying them as the most highly exposed and vulnerable subpopulation (Table 4). Consequently, the toddler model was selected as the worst-case scenario.

**Figure 5:**
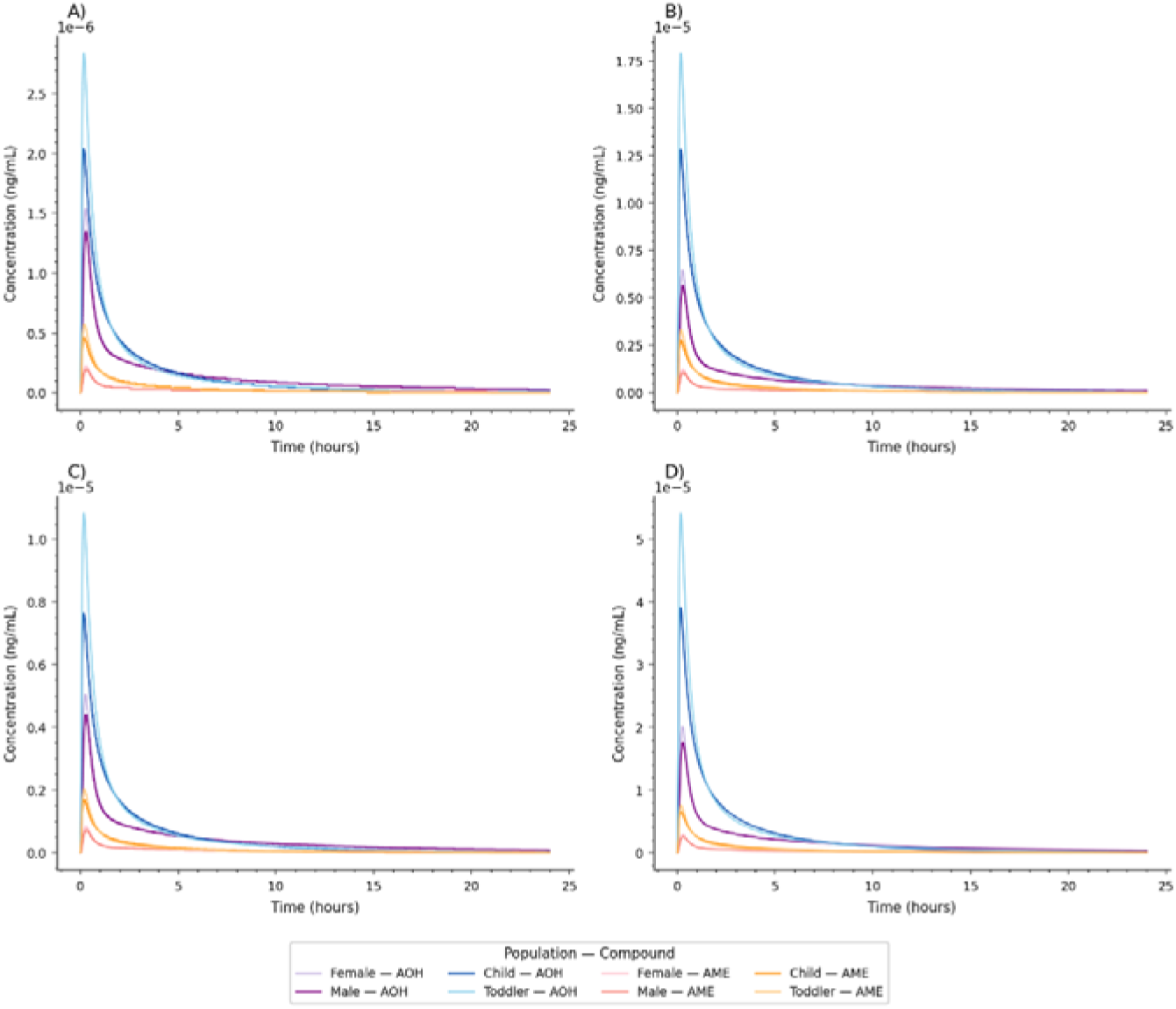
Solid curved depict predicted blood concentration-time profiles for AOH and AME across four human demographic groups. The simulations represent systemic internal exposure to AOH (respectively AME) in adult males (dark purple, respectively dark pink), adult females (light purple, respectively light pink), male children (dark blue, respectively dark yellow), and male toddlers (light blue, respectively light yellow) under various dietary exposure scenarios. (A) Mean exposure low value scenario. (B) Mean exposure high value scenario. (C) 95^th^-percentile low value exposure scenario. (D) 95^th^-percentile high value exposure scenario.

**Table 4:** Predicted blood concentrations of AOH and AME in humans. The numbers next to the toxin represent the scenario number.

| Toxin | Scenario | | | $C_{\max \text{ blood}}$<br>($\times 10^{-5}$ ng/mL) | Toxin | Scenario | | | $C_{\max \text{ blood}}$<br>($\times 10^{-5}$ ng/mL) |
| --- | --- | --- | --- | --- | --- | --- | --- | --- | --- |
| AOH | 2 | Mean | Low | 0.14 | AME | 6 | Mean | Low | 0.02 |
|  |  |  | High | 0.56 |  |  |  | High | 0.10 |
|  |  | 95<sup>th</sup><br>percentile | Low | 0.44 |  |  | 95<sup>th</sup><br>percentile | Low | 0.07 |
|  |  |  | High | 1.76 |  |  |  | High | 0.25 |
|  | 3 | Mean | Low | 0.15 |  | 7 | Mean | Low | 0.02 |
|  |  |  | High | 0.65 |  |  |  | High | 0.12 |
|  |  | 95<sup>th</sup><br>percentile | Low | 0.51 |  |  | 95<sup>th</sup><br>percentile | Low | 0.08 |
|  |  |  | High | 2.01 |  |  |  | High | 0.29 |
|  | 4 | Mean | Low | 0.20 |  | 8 | Mean | Low | 0.05 |
|  |  |  | High | 1.28 |  |  |  | High | 0.28 |
|  |  | 95<sup>th</sup><br>percentile | Low | 0.77 |  |  | 95<sup>th</sup><br>percentile | Low | 0.17 |
|  |  |  | High | 3.90 |  |  |  | High | 0.66 |
|  | 5 | Mean | Low | 0.28 |  | 9 | Mean | Low | 0.06 |
|  |  |  | High | 1.79 |  |  |  | High | 0.33 |
|  |  | 95<sup>th</sup><br>percentile | Low | 1.08 |  |  | 95<sup>th</sup><br>percentile | Low | 0.20 |
|  |  |  | High | 5.41 |  |  |  | High | 0.76 |

### 3.5 Sensitivity analysis and Monte-Carlo simulations

To identify the primary drivers of internal exposure, local sensitivity analysis was performed to quantify the impact of specific parameters on the predicted C_max_ of AOH and AME in systemic circulation of the toddler model (Figure 6).

**Figure 6:**
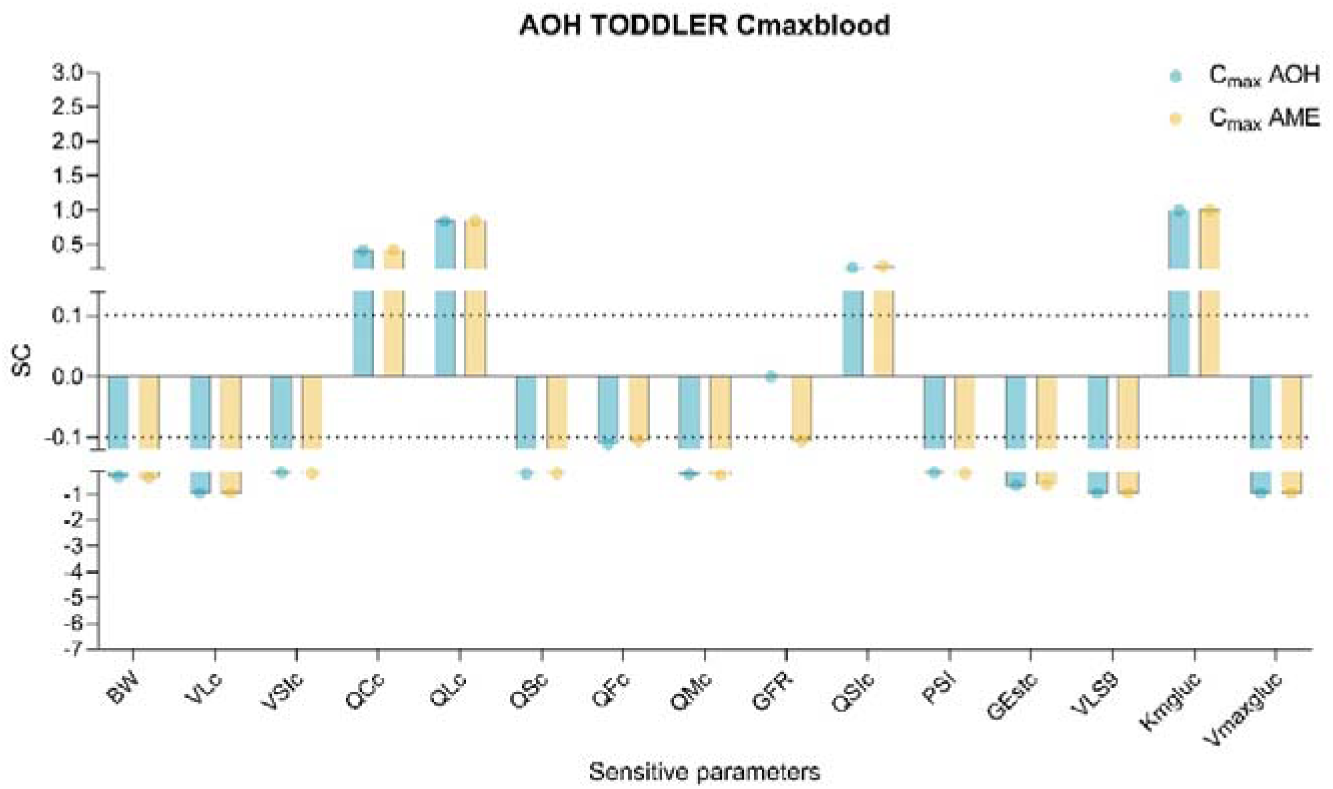
Normalized sensitivity coefficients of predicted peak blood concentrations (Cmax) for the toddler worst-case exposure scenario. Bars represent the normalized sensitivity coefficients for AOH (blue) and AME (yellow). Parameters with coefficients >|0.1| are shown. Abbreviations of the parameters can be found in Table S2.

Parameters describing uptake across the gastrointestinal barrier (VSIc, QSIc, PSI, Gestc) and hepatic biotransformation (VLS9, Kmgluc, Vmaxgluc) appeared to be the most sensitive in the toddler models (Figure 6). The models also exhibited high sensitivity to physiological scaling factors, including body weight and the fractional distribution of blood flow to both highly perfused organs (liver, small intestine) and storage tissues (fat, muscle, slowly perfused tissue).

To further account for inter-individual variability and parameter uncertainty, Monte Carlo simulations (n = 500) were performed for the toddler models. This probabilistic approach integrated the variability of the most sensitive identified parameters to predict the distribution of systemic concentrations across a simulated population. First, a single-exposure scenario was modeled over 24 hours to characterize the acute kinetic profile (Figure 7). Maximal concentrations of AOH and AME reached 5.3*10^-5^ ng/mL and 0.98*10^-5^ ng/mL respectively. As human dietary patterns typically involve three-meal intake throughout the day, a single-dose model may overlook the potential for systemic accumulation. To better reflect realistic exposure, a chronic meal-time scenario was simulated over a 5-day period with dosing events occurring at 08:00 a.m., 12:00 a.m., and 07:00 p.m. (Figure S5). Even at the end of the 5-day period, the peak concentrations for both toxins remained within the same order of magnitude as the single-dose scenario, with respective values of 5.80*10^-5^ ng/mL and 1.05*10^-5^ ng/mL, suggesting that AOH and AME do not accumulate under normal physiological conditions.

**Figure 7:**
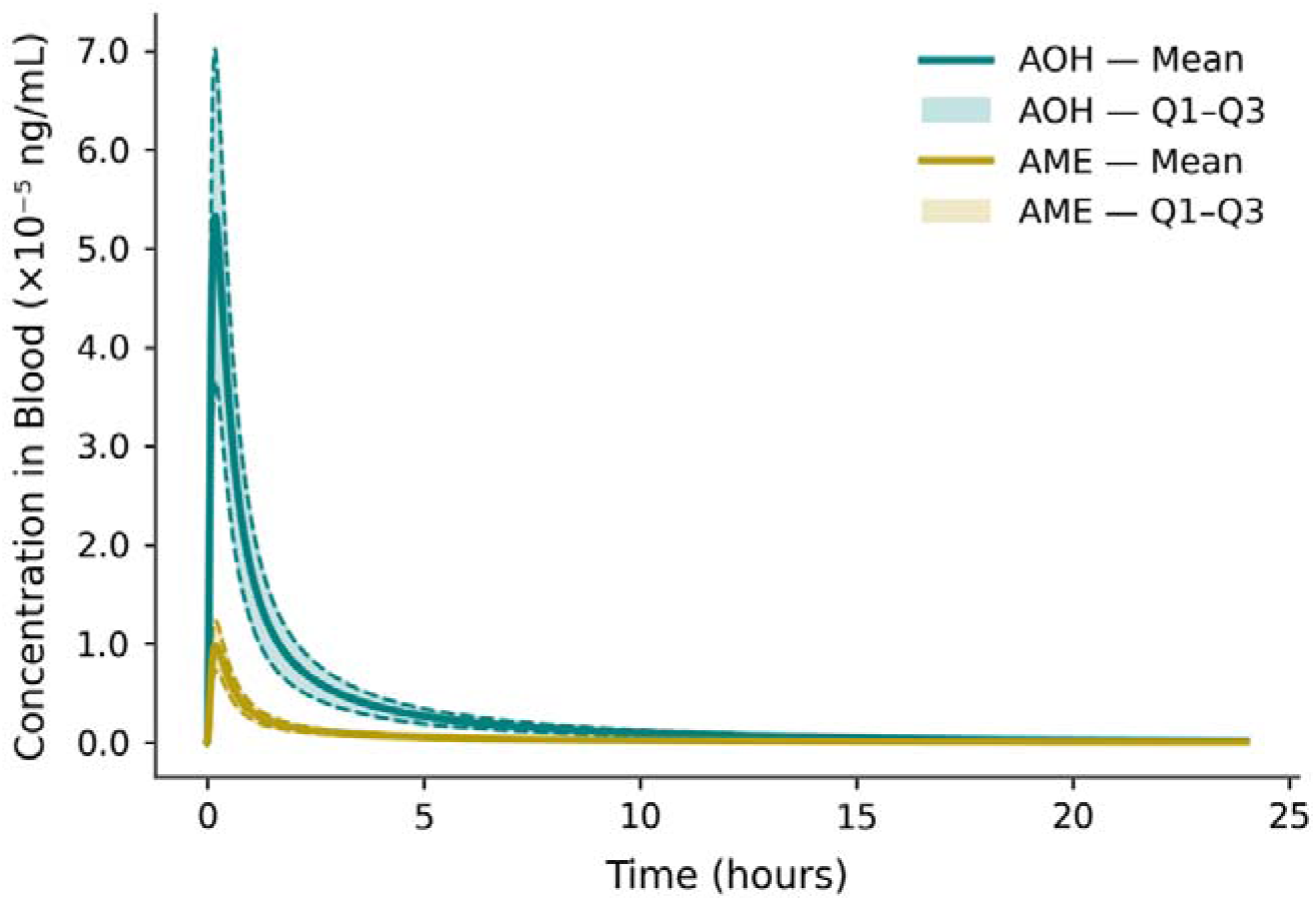
Predicted 24 h acute systemic kinetics of AOH (blue) and AME (yellow) in toddlers. Profiles represent a 95^th^ percentile worst-case scenario (n=500 Monte Carlo iterations) following a single oral exposure. Solid lines indicate the population mean; shaded areas characterize the interquartile distribution (Q1-Q3).

### 3.6 Risk evaluation

While the Monte Carlo simulations provided a robust characterization of the uncertainty and variability of the population for systemic blood concentrations, we further extended our analysis to predict the acute distribution of AOH and AME within specific target organs, further compared with *in vitro* LOAEL values collected from the literature (EFSA 2016). Concentration in the liver, large intestine, kidneys, adipose tissue (assimilated as mammary tissue for female model) and uterus tissue for female model were predicted for AOH and AME following the different exposure scenarios. Models were simulated for acute exposures of 24 h, 48 h and 72 h to match the *in vitro* LOAEL measurement conditions. Values for AME LOAEL upon 72 h acute exposure were not reported and no risk evaluation was evaluated for this condition. Tables of values (Table S6) as well as 48 h (AOH, AME) and 72 h (AOH) figures (Figure S6) are provided in Supplementary information 1.

Across all demographics and exposure scenarios, the predicted organ-specific concentrations did not exceed 10^-4^ µM. In contrast, the lowest LOAEL value was not below the micromolar range for 24 h acute exposure scenario (Figure 8).

**Figure 8:**
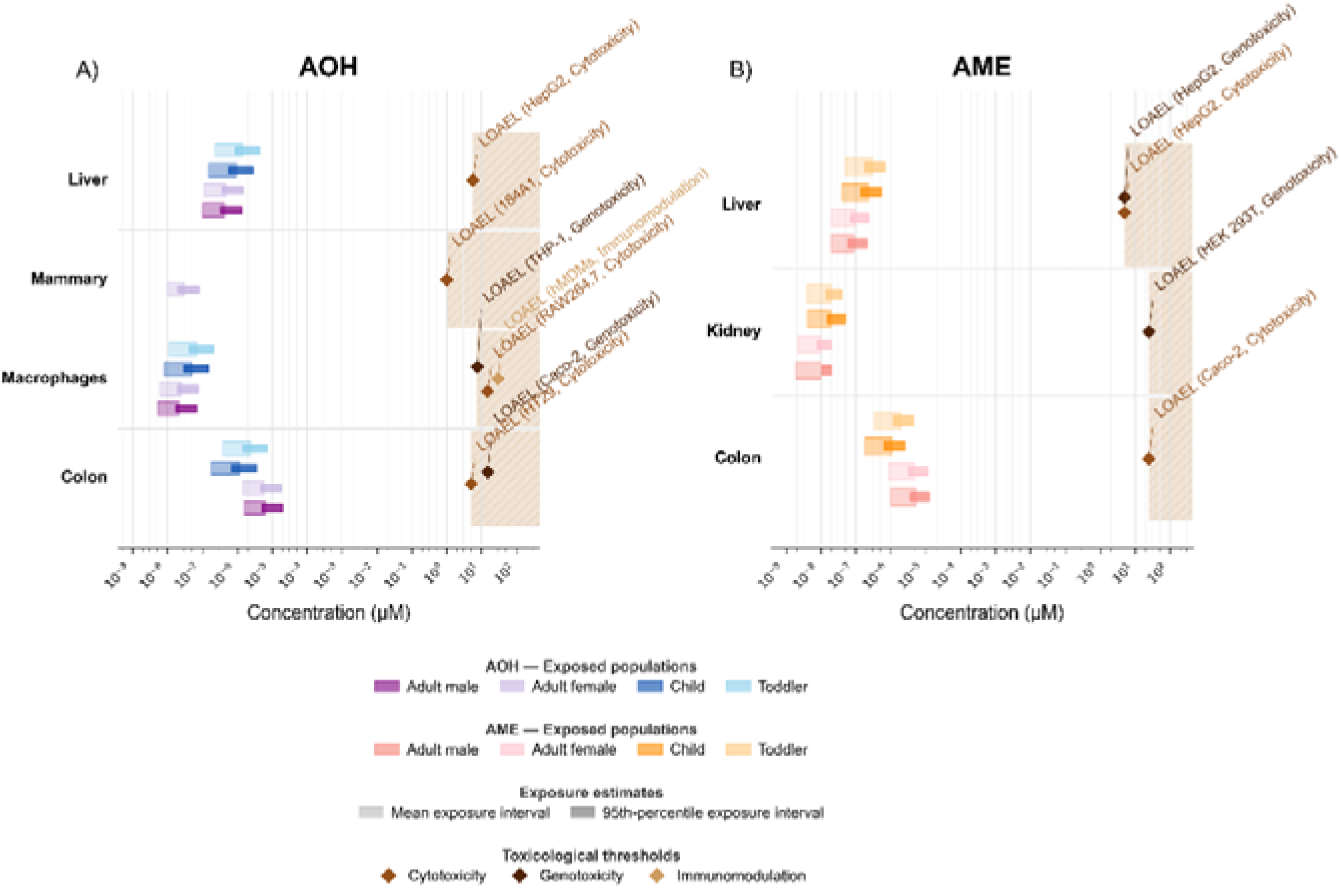
Comparative risk evaluation of tissue-specific internal doses and *in vitro* LOAELs in females, males, male children and male toddlers. (A) The plot illustrates the distribution of AOH in the liver, kidney, colon, and mammary-representative adipose tissue compared to established toxicological thresholds after 24 h acute exposure. (B) The plot illustrates the distribution of AME in the liver, kidney and colon compared to established toxicological thresholds after 24 h acute exposure.

Comparing these values revealed a significant MoS of at least four orders of magnitude. Even for the toddler model, which was identified as the most highly exposed subpopulation and worst-case scenario, the internal C_max_ values are approximately 10,000-fold lower than the concentrations required to induce a toxicological response *in vitro*. Similarly, the MoS for the endocrine disruption endpoint in the uterus tissue (Figure S6) reached the 10^8^ order of magnitude. This significant gap suggests that under current dietary exposure scenarios defined by the EFSA (2016) report, AOH and AME are unlikely to reach systemic or tissue-specific levels that would pose an acute human health concern, at least regarding single toxin exposure.

Despite the substantial MoS identified, several uncertainties inherent to the QIVIVE framework warrant consideration. First, nominal concentrations used to establish *in vitro* LOAELs may not accurately reflect free intracellular concentrations due to plastic adsorption to assay plates or non-specific binding to serum proteins, potentially leading to overestimation of intracellular concentrations of AOH and AME. A better characterization of actual exposure and the fate of chemicals within the cellular test system would thus increase QIVIVE accuracy. A recently published method to account for such effects (Brenner *et al*. 2026) was unfortunately not applicable due to the required extensive reporting of methodical specifications in toxicity testing that were not met by most of the included studies.

Second, while toxicity was reported for AOH and AME as isolated toxins, real-world exposure involves complex mycotoxin mixtures of a multitude of chemicals with differing structures and bioactivities, that have previously been reported to cause synergistic toxicity at low concentrations (Vejdovszky *et al*. 2017a, b). Additive or synergistic interactions may occur even at concentrations below individual LOAELs, but are not considered in single-compound risk evaluation. Finally, while AOH and AME were prioritized based on their high prevalence in the food supply, other emerging toxins from the *Alternaria* metabolome exhibit significantly higher toxicity potency, particularly concerning genotoxicity (Schwarz *et al*. 2012; Aichinger *et al*. 2022) and may contribute disproportionately to overall risk despite lower exposure levels. Consequently, comprehensive cumulative risk assessment incorporating mixture effects and a hazard characterization of the broader *Alternaria* spp. metabolome should be prioritized in future research.

## 4 Conclusion

This study successfully integrated *in vitro* glucuronidation kinetics with *in silico* PBK modeling to provide refined risk evaluation of the single *Alternaria* toxins AOH and AME. Our results establish that glucuronidation serves as a highly efficient clearance pathway for both toxins in both rodents and humans, with V_max_ and K_m_ merging as sensitive parameters in the PBK model. Interspecies scaling and probabilistic Monte Carlo simulations revealed that even though toddlers represent the most vulnerable subpopulation under worst-case scenarios, systemic accumulation is unlikely under realistic chronic dietary exposure. Notably, predicted internal doses remained more than four orders of magnitude below *in vitro* LOAELs, establishing a MoS exceeding 10,000.

While these findings indicate that AOH and AME are unlikely to pose acute human health risk at current dietary exposure levels, several uncertainties remain. Updated exposure estimations could directly be implemented in the model to update the risk evaluation. The model does not account for potential enterohepatic recirculation, which could prolong systemic exposure if biliary-excreted glucuronides undergo intestinal deconjugation and reabsorption. Moreover, nominal *in vitro* concentrations may overestimate bioactive free intracellular levels due to non-specific binding to plasticware or serum protein considerations. As discussed, cross-species application of rat-derived glucuronidation parameters to the mouse model introduces quantitative uncertainty. Finally, this study focused on isolated AOH and AME, while dietary exposure involves complex mycotoxin mixtures. As other *Alternaria* metabolites may contribute disproportionately to overall risk through additive or synergistic effects, these limitations warrant further investigations. Nevertheless, this work establishes a robust computational framework to close the data gap of toxicokinetics for risk assessment of the single mycotoxins AOH and AME exposure.

## Associated content

### Supporting information

The Supporting Information is available free of charge at “xxx to be inserted at a later stage xxx”.

### Author Contributions

All authors have approved the final version of the manuscript.

CRediT:

E.B., B.D., E.V., D.M., G.A. conceptualization; E.B., B.D., N.H. data curation, formal analysis, investigation, validation, visualization; E.B., B.D. writing – original draft; E.V., A.G. methodology, E.V., D.M., G.A. supervision; E.D., A.G. project administration; D.M., A.G. resources, funding acquisition; all authors writing – review & editing.

## Funding

The European Partnership for the Assessment of Risks from Chemicals has received funding from the European Union’s Horizon Europe research and innovation program under Grant Agreement No 101057014 and has received co-funding of the authors’ institutions. Views and opinions expressed are, however, those of the author(s) only and do not necessarily reflect those of the European Union or the Health and Digital Executive Agency. Neither the European Union nor the granting authority can be held responsible for them. Open access funding was provided by the University of Vienna.

## Notes

The authors declare no competing financial interest.

## Acknowledgements

This research was supported using resources of the VetCore Facility (Mass Spectrometry) of the University of Veterinary Medicine Vienna. The authors sincerely thank Jelena Vukoje and Vice Tomičić for conducting exploratory incubation experiments with rat liver S9 fractions and self-prepared buffer solutions. Figure 1 was created in BioRender. Aichinger, G. (2026) https://BioRender.com/aziuwd2.

## Notes

### Competing Interest Statement

The authors have declared no competing interest.

### Summary of Updates

Figure duplications caused by PDF conversion were removed. The revision concerns formatting issues only; no scientific content was changed.

