## Supplementary material for "Human internal exposures to alternariol and its monomethyl ether are predicted below thresholds of *in vitro* toxicity by physiologically based kinetic modeling": S1 Supplementary files.pdf

#### 1 Materials and Methods

##### 1.1 Glucuronidation of AOH and AME in rat and human liver S9 fractions

**Chemicals and biological materials – synthesis of AME.** The convergent synthesis of AME was carried out according to Mikula *et al.* (2013). The building blocks were prepared from orcinol and 3,5-dimethoxyaniline respectively and coupled using diethyl chlorophosphate (DECP). The benzocoumarin core structure was established by intramolecular cyclization of the iodoresorcylic acid phenyl ester. Selective demethylation of alternariol 2,4-dimethyl ether (ADME) with $AlCl_3$/NaI yielded AME which was isolated using reverse phase column chromatography. The identity of the material was confirmed by analytical HPLC-ESI-MS(MS) and $^1H$ NMR spectroscopy.

**LC-MS/MS analysis of the parent toxins and their glucuronide metabolites.** Quantification of the parent compounds and semi-quantification of their glucuronide metabolites were performed on an Agilent 1290 Infinity II high-performance liquid chromatography system (Agilent Technologies, Santa Clara, CA, USA) coupled to a SCIEX QTrap 6500+ (Malborough, MA, USA) mass spectrometer, equipped with an electrospray ionization interface.

Chromatographic separation was achieved using an Ascentis Express C18 column (10 cm × 2.1 mm, 2.7 µm, Supelco, Munich, DE), equipped with a Phenomenex SecurityGuard™ C18 cartridge (4 × 2.0 mm ID, Phenomenex Ltd. Deutschland, Aschaffenburg, Germany). Eluent A was aqueous ammonium acetate (5 mM, pH adjusted to 8.6 with a 25% ammonia) and eluent B was methanol. During the first minute of the gradient elution, the column was kept at 10% eluent B. Subsequently, the methanol content was linearly raised to 38% by 5.5 min. Then, the percentage of eluent B was raised to 60% at 7 min and 100% at 7.5 min. This condition was then held for 2 min, and finally, the column was re-equilibrated at the initial conditions, resulting in a total run time of 11.5 min, with a constant flow rate of 0.4 mL/min.

The autosampler compartment was cooled to 10 °C, the column oven temperature was kept at 30 °C, and the injection volume was set to 5 µL. The tandem mass spectrometer was operated in multiple reaction monitoring (MRM) mode using negative ionization, detecting the analytes in their deprotonated forms. The following conditions were applied: curtain gas 35, source voltage -4500 V, capillary temperature 450 °C, and ion source gases 1 and 2 set to 60, respectively. Data acquisition and initial evaluation were performed using Analyst (v. 1.7.) and SCIEX OS (v. 3.3.1.43), both developed by SCIEX LLC (Marlborough, MA, USA). Further data processing was conducted using the software Skyline (version 25.1., MacCoss Lab, Department of Genome Sciences, University of Washington, Seattle, WA, USA).

While the MS parameters for AOH/AME could be automatically optimized, method optimization of their metabolites relied on repeated injections of selected incubation samples containing the peaks of interest due to the absence of reference standards. The samples were injected via the LC autosampler, and relevant MS parameters, including declustering potential (DP) and collision energy (CE), were manually optimized to enhance signal intensity. The MRM transitions and corresponding semi-optimized MS parameters are summarized in Table S1. In the samples, the parent toxins AOH and AME were quantified via solvent-based external calibration of certified analytical standards, while their metabolites were semi-quantified due to the lack of commercially available standards.

### 2 Tables and Figures

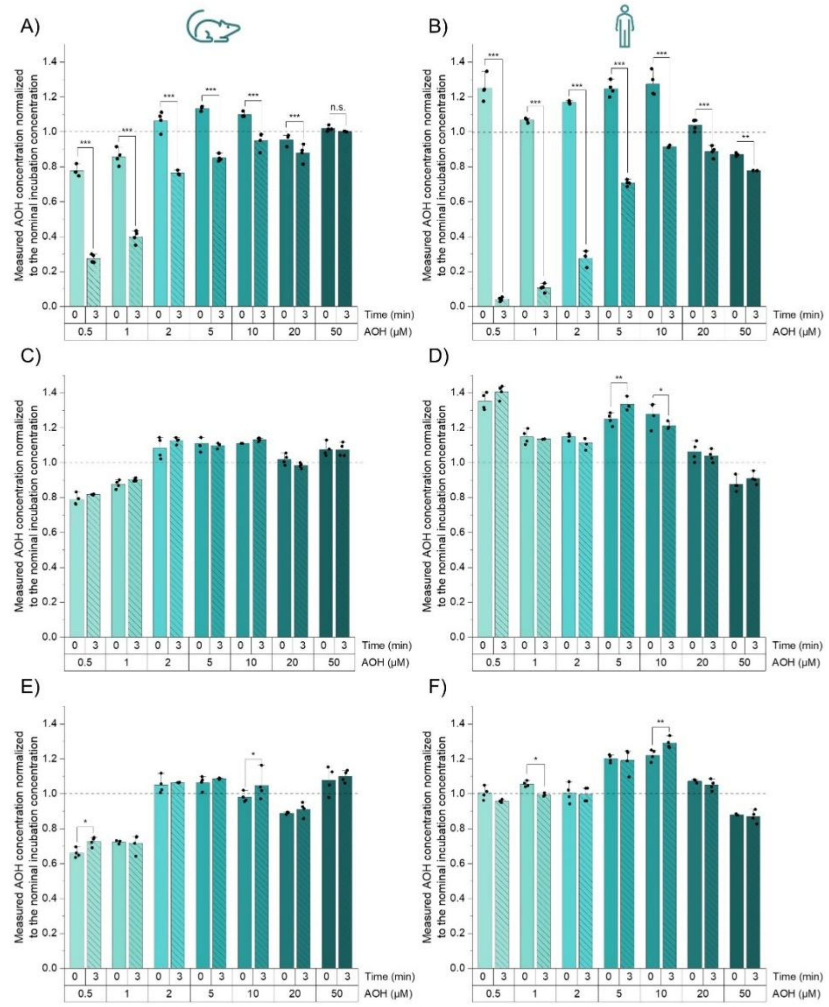


**Figure S1:** Parent compound levels after the incubation of rat (A, C, E) and human (B, D, F) liver S9 fractions with 0.5-50 µM AOH for 3 minutes. Figures A and B show the standard incubation protocols, C and D refer to UDGPA-free indications, and E and F show samples additionally incubated with β-glucuronidase. Columns represent the mean ± SD of four independent experiments. Normality was assessed (Shapiro–Wilk), and outliers were evaluated using the Nalimov test and excluded where appropriate. One-way ANOVA, followed by Fisher's LSD posthoc test, was used to detect significant differences. Significance is indicated as follows: n.s. → not significant; * → $0.01 < p < 0.05$; ** → $0.001 < p < 0.01$; and *** → $p < 0.001$. If not indicated otherwise, the difference between 0 and 3 minutes within the same incubation condition was not statistically significant.

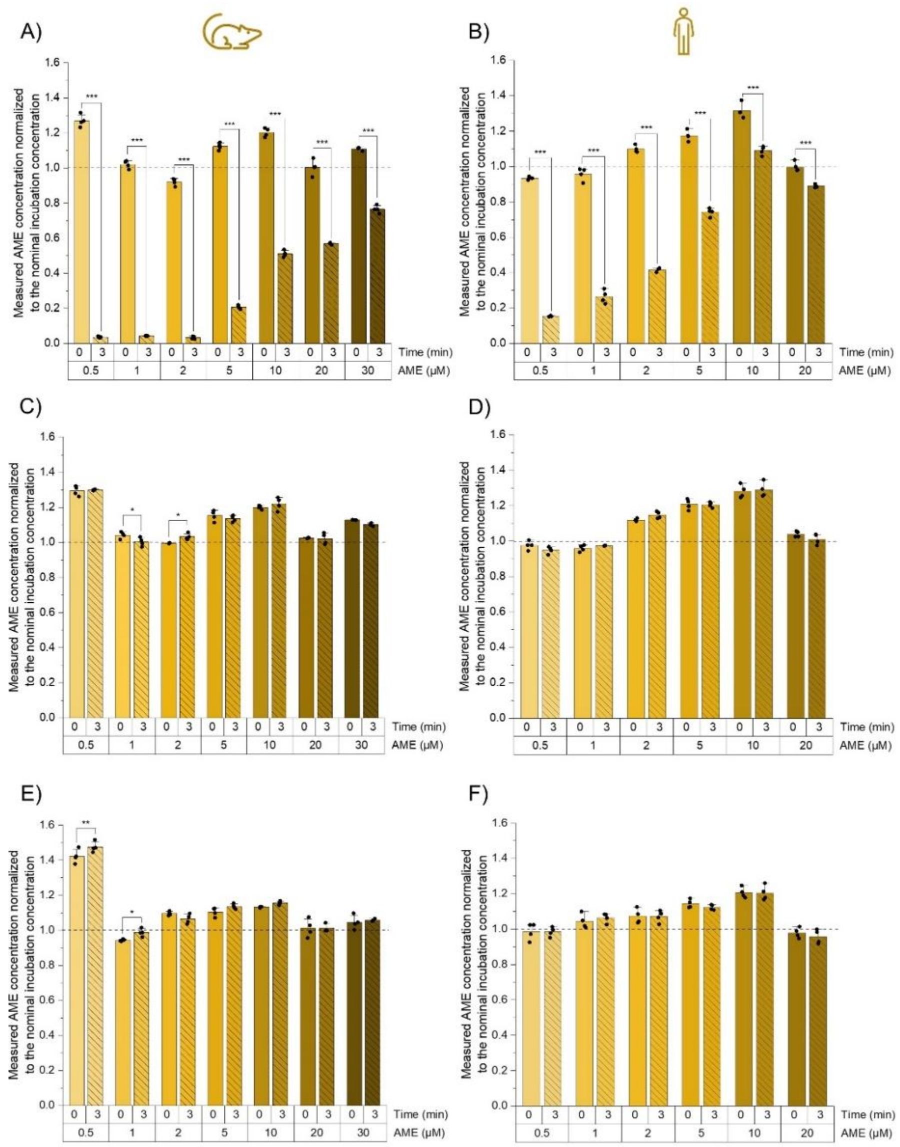


**Figure S2:** Parent compound levels after the incubation of rat (A, C, E) and human (B, D, F) liver S9 fractions with 0.5-30 µM (rat) and 0.5-20 µM (human) AME for 3 minutes. Figures A and B show the standard incubation protocols, C and D refer to UDGPA-free indications, and E and F show samples additionally incubated with β-glucuronidase. Columns represent the mean ± SD of four independent experiments. Normality was assessed (Shapiro–Wilk), and outliers were evaluated using the Nalimov test and excluded where appropriate. One-way ANOVA, followed by Fisher's LSD posthoc test, was used to detect significant differences. Significance is indicated as follows: * → $0.01 < p < 0.05$; ** → $0.001 < p < 0.01$; and *** → $p < 0.001$. If no asterisk is shown, the difference between 0 and 3 minutes within the same incubation condition was not statistically significant.

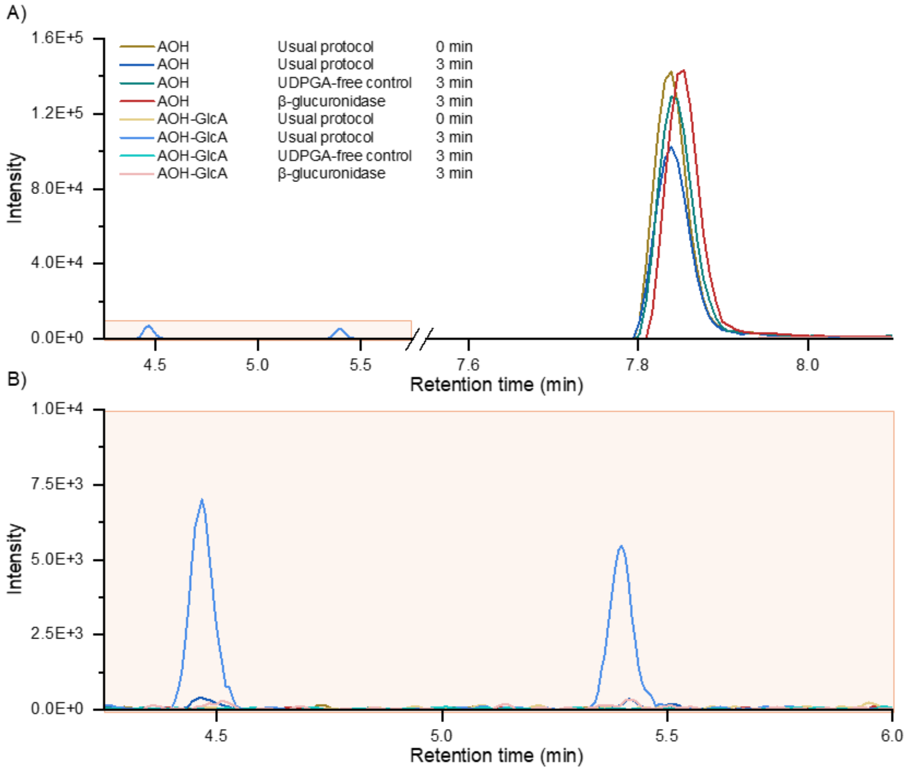


**Figure S3:** Exemplary chromatograms of AOH (A) and its glucuronide metabolites (B) in rat liver S9 fractions after incubation with 20 µM AOH. Each line depicts the MRM transition of the highest intensity of AOH or AOH-GlcA from one representative replicate of each condition.

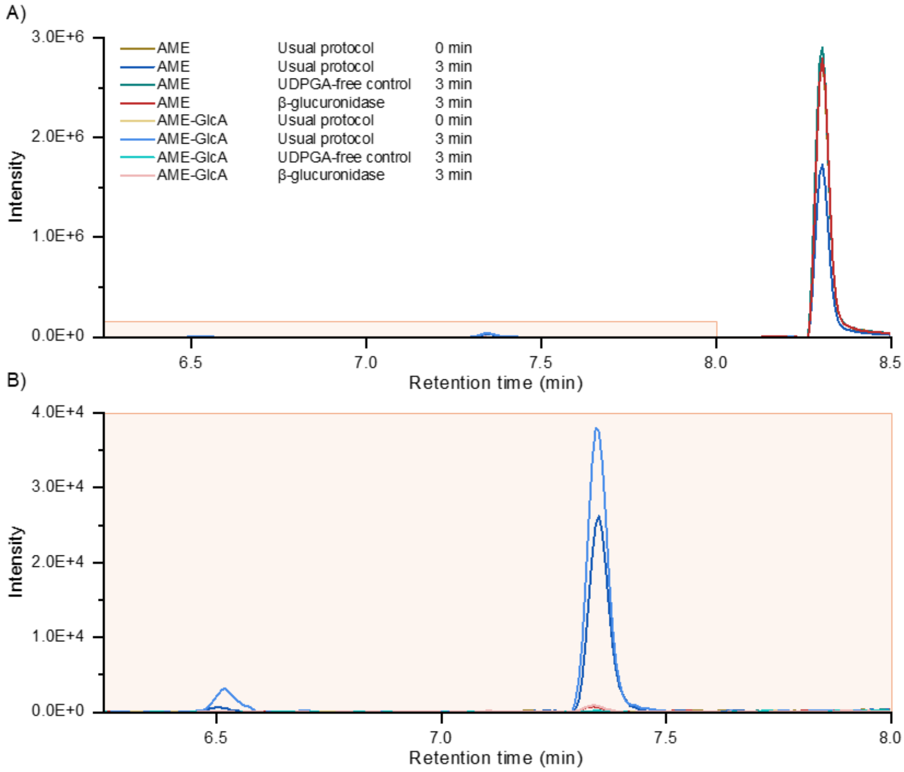


**Figure S4:** Exemplary chromatograms of AME (A) and its glucuronide metabolites (B) in rat liver S9 fractions after incubation with 20 µM AME. Each line depicts the MRM transition of the highest intensity of AME or AME-GlcA from one representative replicate of each condition. At the retention times of the AME-GlcA, in source fragmentation to AME is visible.

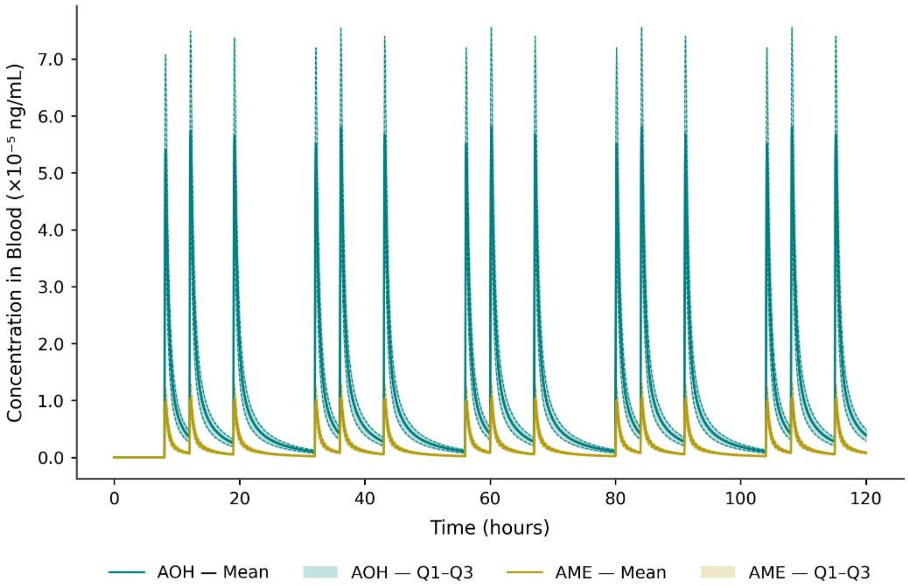


**Figure S5:** Simulated 5-day chronic dietary exposure of AOH (blue) and AME (yellow) in toddlers. Profiles represent a 95$^{th}$ percentile worst-case scenario (n = 500 Monte Carlo iterations) with three daily intake events. Solid lines depict the population mean and shaded areas characterize interquartile distribution.

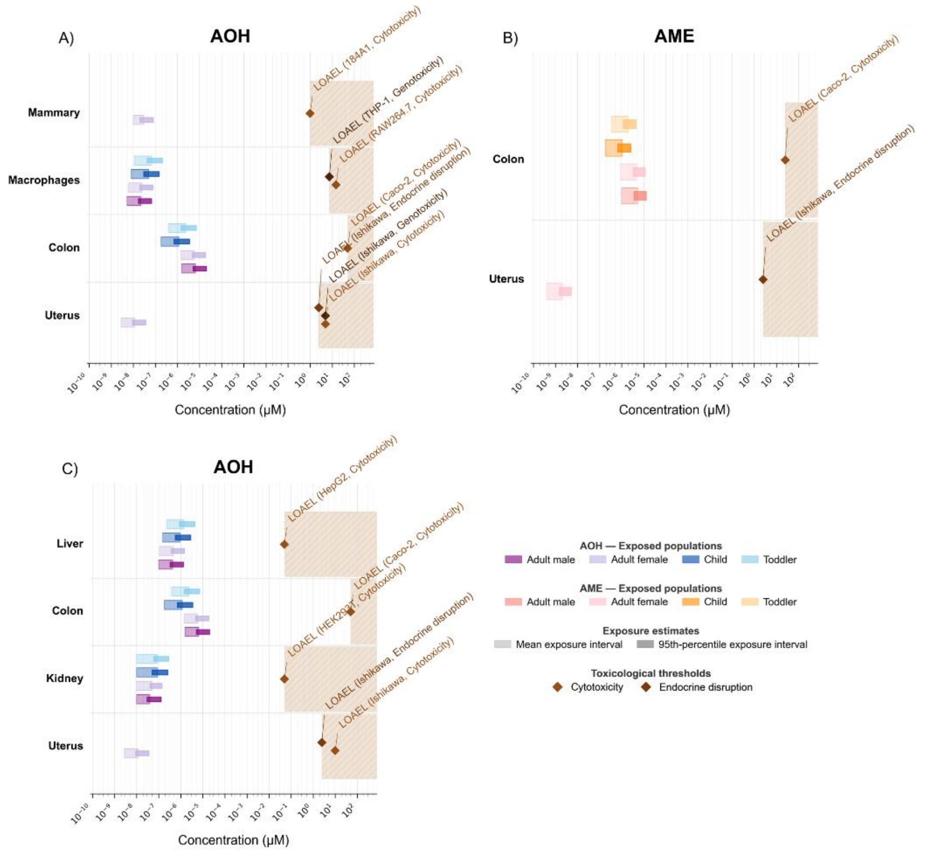


**Figure S6:** Comparative risk evaluation of tissue-specific internal doses and *in vitro* lowest observed adverse effect levels (LOAELs) in females, males, male children and male toddlers. (A) The plot illustrates the distribution of AOH in the mammary-representative adipose tissue, macrophages, colon and uterus tissue compared to established toxicological thresholds after 48 h acute exposure. (B) The plot illustrates the distribution of AME in the colon and uterus tissue compared to established toxicological thresholds after 48 h acute exposure. (C) The plot illustrates the distribution of AOH in the liver, colon, kidney and uterus tissue compared to established toxicological thresholds after 72 h acute exposure.

**Table S1.** Mass spectrometric parameters of the analytes measured via LC-MS/MS in negative ionization mode. DP – declustering potential, CE – collision energy; CXP – collision cell exit potential. An entrance potential of -10 V and a dwell time of 10 ms was universally set. Quantifier MRM transitions are marked bold where applicable.

| Molecule name | Precursor *m*/*z* | Product *m*/*z* | DP (V) | CE (V) | CXP (V) |
|---|---|---|---|---|---|
| Alternariol (AOH) | 256.9 | **212.9** | **-120** | **-32** | **-15** |
| | | 214.9 | -120 | -36 | -13 |
| | | 211.9 | -120 | -40 | -13 |
| Alternariol monomethyl ether (AME) | 270.9 | 227.9 | -125 | -40 | -15 |
| | | **255.9** | **-125** | **-30** | **-17** |
| | | 182.9 | -125 | -54 | -13 |
| Alternariol-glucuronide (AOH-GlcA) | 433.1 | 257.0 | -40 | -40 | -11 |
| | | 215.0 | -40 | -65 | -11 |
| | | 175.0 | -40 | -20 | -11 |
| Alternariol monomethyl ether-glucuronide (AME-GlcA) | 447.1 | 271.1 | -20 | -30 | -11 |
| | | 256.0 | -20 | -60 | -11 |

**Table S2**: Physiological parameters. a – similar values as male values.

| | **Mouse** | **Man** | **Woman** | **Male child** | **Male toddler** |
|---|---|---|---|---|---|
| Body weight | 0.037 | 73 | 60 | 19 | 10 |
| **Relative Organ Weight (Percent Body Weight)** | | | | | |
| VLc – Liver fraction | 5.49 | 3.19 | 2.98 | 3.74 | 3.80 |
| VPc- Portal vein fraction | 1.55 | 0.63 | 0.64 | 0.68 | 0.56 |
| VRc – Rapidly perfused tissue fraction | 1.23 | 2.12 | 1.99 | 2.10 | 1.98 |
| VSc – Slowly perfused tissue fraction | 27.26 | 19.63 | 17.48 | 16.53 | 15.70 |
| VFc – Adipose tissue fraction | 0.48 | 20.36 | 30.55 | 19.32 | 23.43 |
| VMc – Muscles tissue fraction | 38.40 | 40.74 | 29.85 | 30.51 | 19.53 |
| VKc – Kidney tissue fraction | 1.67 | 0.57 | 0.59 | 0.73 | 0.80 |
| VSIc – Small intestine tissue fraction | 2.53 | 1.17 | 1.25 | 1.44 | 1.04 |
| VLIc – Large intestine tissue fraction | 1.09 | 0.67 | 0.74 | 0.79 | 0.61 |
| VBc – Blood fraction | - | 7.26 | 6.50 | 7.37 | 5.00 |
| VUc – Uterus fraction | - | - | 0.16 | - | - |
| **Volumes (L) and surface areas ($dm^2$)** | | | | | |
| VB – Volume of blood | 0.08 *bw | - | - | - | - |
| VB – Volume of blood | 0.08 *bw | - | - | - | - |
| VSIL – Volume of small intestine lumen | $2.85*10^{-3}$ | 4.45 | 4.13 | 0.48 | 0.34 |
| VLIL – Volume of large intestine lumen | $1.27*10^{-3}$ | 2.38 | 2.16 | 1.19 | 0.23 |
| areaSI – Area of the small intestine lumen | 0.36 | 2917.50 | 2709.07 | 1771.31 | 1250.34 |
| areaLI – Area of the large intestine lumen | 0.12 | 107.80 | 98.02 | 73.51 | 58.81 |

**Table S2 *continued*:** Physiological parameters. a – similar values as male values

| | Mouse | Man | Woman | Male child | Male toddler |
|---|---|---|---|---|---|
| **Blood flow rates (Percent Cardiac Output)** | | | | | |
| QCc – Cardiac output (L/min) | 0.275*bw ^0.75 | 6.5 | 5.9 | 3.4 | 1.2 |
| QPc – Portal vein | 14.1 | 5.0 | a | a | a |
| QLc – Liver | 16.2 | 6.5 | a | a | a |
| QRc – Rapidly perfused tissues | 7.1 | 104.0 | 105.0 | a | a |
| QSc – Slowly perfused tissues | 18.0 | 10.0 | a | a | a |
| QFc – Adipose tissue | 7.0 | 5.0 | 8.5 | a | a |
| QMc – Muscles tissues | 15.9 | 17.0 | 12.0 | a | a |
| QKc – Kidney tissues | 9.1 | 19.0 | 17.0 | a | a |
| QSIc – Small intestine tissues | 13.5 | 10.0 | 5.0 | a | |
| QUc – Uterus tissues | - | - | 0.4 | - | - |
| **Further physiological parameters** | | | | | |
| GFR – Glomerular filtration rate (mL/min) | 0.28 | 125.00 | a | a | a |
| GEstc – Time to empty 50% of stomach (min) | 74.0 | 11.8 | a | a | a |
| tpassSI – Transit time small intestine | 1.5 | 4.0 | a | a | a |
| tpassLI – Transit time large intestine | 8 | 36 | 48 | 34 | 32 |

**Table S3:** Physicochemical parameters

| | **Physicochemical properties** | | | | | **Permeability coefficient (logcm/s)** | | | |
|---|---|---|---|---|---|---|---|---|---|
| | MW – Molar mass (g/mol) | pKa1 | pKa2 | LogP | LogD | Kamiya model | Lanevskij model | *In vitro* | Extrapolation |
| AOH | 258.23 | 7.63 | 8.44 | 3.18 | 3.17 | -4.57 | -4.38 | -5.09 | - |
| AME | 272.25 | 7.71 | - | 3.32 | 3.32 | -4.60 | -4.39 | - | -5.11 |

**Table S4:** Blood/Tissue partitioning (QIVIVE). a – similar values as male values; (A) – AOH; (B) – AME

(A)

| AOH | Mouse | Man | Woman | Male child | Male toddler |
|---|---|---|---|---|---|
| PL – Liver | 2.2 | 3.21 | a | a | a |
| PR – Rapidly perfused | 2.215 | 0.893 | 0.894 | 0.898 | 0.920 |
| PS – Slowly perfused | 2.621 | 4.601 | 4.635 | 4.733 | 4.627 |
| PF – Adipose tissues | 7.9 | 7.3 | a | a | a |
| PM – Muscles tissues | 1.39 | 2.09 | a | a | a |
| PK – Kidney tissues | 2.06 | 2.07 | a | a | a |
| PSI – Small intestine tissue | 3.07 | 3.97 | a | a | a |
| FU – Fraction unbound | 0.06 | 0.06 | a | a | a |

(B)

| AME | Man | Woman | Male child | Male toddler |
|---|---|---|---|---|
| PL – Liver | 3.8 | a | a | a |
| PR – Rapidly perfused | 0.982 | 0.982 | 0.988 | 1.014 |
| PS – Slowly perfused | 5.509 | 5.551 | 5.560 | 5.540 |
| PF – Adipose tissues | 7.1 | a | a | a |
| PM – Muscles tissues | 2.44 | a | a | a |
| PK – Kidney tissues | 2.41 | a | a | a |
| PSI – Small intestine tissue | 4.72 | a | a | a |
| FU – Fraction unbound | 0.058 | a | a | a |

**Table S5:** Coefficients of variation (CV) and distribution of sensitive parameters.

| Parameter | CV (%) | Distribution | Reference |
|---|---|---|---|
| BW | 15 | Lognormal | (Price *et al.* 2003) |
| VLc | 30 | Normal | (Clewell and Clewell 2008) |
| VSIc | 30 | Normal | (Clewell and Clewell 2008) |
| QCc | 30 | Normal | (Clewell and Clewell 2008) |
| QLc | 30 | Normal | (Clewell and Clewell 2008) |
| QSc | 30 | Normal | (Clewell and Clewell 2008) |
| QFc | 30 | Normal | (Clewell and Clewell 2008) |
| QMc | 30 | Normal | (Clewell and Clewell 2008) |
| QSIc | 30 | Normal | (Clewell and Clewell 2008) |
| GFR | 18 | Lognormal | (Fravel *et al.* 2023) |
| PSI | 30 | Lognormal | (Clewell *et al.* 1999) |
| Gestc | 69 | Lognormal | (Oberle *et al.* 1990) |
| VLS9 | 26 | Lognormal | (Bhatt *et al.* 2019) |
| KmglucAOH | 31.96 | Lognormal | Estimated |
| VmaxglucAOH | 9.64 | Lognormal | Estimated |
| KmglucAME | 65.98 | Lognormal | Estimated |
| VmaxglucAME | 8.57 | Lognormal | Estimated |

**Table S6:** Prediction values from the comparative risk evaluation of tissue-specific internal doses and *in vitro* lowest observed adverse effect levels (LOAELs) in females, males, male children and male toddlers after 24 h, 48 h and 72 h acute exposure of AOH and AME. (A) Low mean acute exposure scenario dose. (B) High mean acute exposure scenario dose. (C) Low 95th percentile acute exposure scenario dose. (D) High 95th percentile acute exposure scenario dose. AUC – Area under the curve.

(A)

| Compound | Population | Compartment | $C_{max}$ (µM) | $T_{max}$ | AUC |
|---|---|---|---|---|---|
| AOH | Male | Blood | 5.23E-09 | 0.2768 | 3.04E-06 |
| | | Large Intestine Tissue | 1.56E-06 | 0.42502 | 1.95E-06 |
| | | Liver | 1.00E-07 | 0.15186 | 5.08E-08 |
| | | Kidney Tissue | 1.00E-08 | 0.12924 | 5.97E-09 |
| | Female | Blood | 5.96E-09 | 0.24432 | 3.20E-06 |
| | | Large Intestine Tissue | 1.42E-06 | 0.44568 | 1.77E-06 |
| | | Liver | 1.10E-07 | 0.14432 | 5.44E-08 |
| | | Adipose Tissue | 1.00E-08 | 0.94418 | 6.63E-08 |
| | | Kidney Tissue | 1.00E-08 | 0.11192 | 6.28E-09 |
| | | Uterus Tissue | 2.80E-09 | - | - |
| | Male child | Blood | 7.90E-09 | 0.16782 | 3.39E-06 |
| | | Large Intestine Tissue | 1.80E-07 | 0.26978 | 1.88E-07 |
| | | Liver | 1.50E-07 | 0.09384 | 6.15E-08 |
| | | Kidney Tissue | 1.00E-08 | 0.05222 | 1.06E-08 |
| | Male toddler | Blood | 1.10E-08 | 0.1708 | 3.79E-06 |
| | | Large Intestine Tissue | 3.80E-07 | 0.16274 | 1.99E-07 |
| | | Liver | 2.30E-07 | 0.10634 | 9.62E-08 |
| | | Kidney Tissue | 1.00E-08 | 0.05234 | 9.12E-09 |

**Table S6 *continued*:**

(A)

| Compound | Population | Compartment | $C_{max}$ (µM) | $T_{max}$ | AUC |
|---|---|---|---|---|---|
| AME | Male | Blood | 6.98E-10 | 0.24628 | 4.61E-07 |
| | | Large Intestine Tissue | 9.80E-07 | 0.4191 | 1.25E-06 |
| | | Liver | 2.00E-08 | 0.15794 | 7.92E-09 |
| | | Kidney Tissue | 1.88E-09 | - | - |
| | Female | Blood | 8.08E-10 | 0.23414 | 4.83E-07 |
| | | Large Intestine Tissue | 8.90E-07 | 0.42452 | 1.14E-06 |
| | | Liver | 2.00E-08 | 0.12608 | 8.67E-09 |
| | | Kidney Tissue | 2.12E-09 | - | - |
| | | Uterus Tissue | 4.03E-10 | - | - |
| | Male child | Blood | 1.69E-09 | 0.164 | 7.90E-07 |
| | | Large Intestine Tissue | 1.80E-07 | 0.28242 | 1.90E-07 |
| | | Liver | 4.00E-08 | 0.08678 | 1.58E-08 |
| | | Kidney Tissue | 4.16E-09 | - | - |
| | Male toddler | Blood | 2.09E-09 | 0.16206 | 7.92E-07 |
| | | Large Intestine Tissue | 3.40E-07 | 0.16138 | 1.80E-07 |
| | | Liver | 5.00E-08 | 0.07794 | 2.22E-08 |
| | | Kidney Tissue | 4.01E-09 | - | - |

**Table S6 *continued*:**

(B)

| Compound | Population | Compartment | $C_{max}$ (µM) | $T_{max}$ | AUC |
|---|---|---|---|---|---|
| AOH | Male | Blood | 2.18E-08 | 0.27728 | 1.27E-05 |
| | | Large Intestine Tissue | 6.54E-06 | 0.45058 | 8.18E-06 |
| | | Liver | 4.30E-07 | 0.19428 | 2.16E-07 |
| | | Kidney Tissue | 4.00E-08 | 0.19938 | 6.41E-08 |
| | Female | Blood | 2.50E-08 | 0.25534 | 1.34E-05 |
| | | Large Intestine Tissue | 5.93E-06 | 0.45746 | 7.43E-06 |
| | | Liver | 4.70E-07 | 0.17272 | 2.31E-07 |
| | | Adipose Tissue | 3.00E-08 | 1.79882 | 3.76E-07 |
| | | Kidney Tissue | 5.00E-08 | 0.23834 | 6.48E-08 |
| | | Uterus Tissue | 1.17E-08 | - | - |
| | Male child | Blood | 4.96E-08 | 0.17214 | 2.14E-05 |
| | | Large Intestine Tissue | 1.16E-06 | 0.3529 | 1.19E-06 |
| | | Liver | 9.40E-07 | 0.11406 | 3.91E-07 |
| | | Kidney Tissue | 9.00E-08 | 0.15854 | 1.27E-07 |
| | Male toddler | Blood | 6.93E-08 | 0.17528 | 2.39E-05 |
| | | Large Intestine Tissue | 2.38E-06 | 0.1694 | 1.26E-06 |
| | | Liver | 1.43E-06 | 0.11 | 6.11E-07 |
| | | Kidney Tissue | 9.00E-08 | 0.12918 | 1.03E-07 |

**Table S6 *continued*:**

(B)

| Compound | Population | Compartment | $C_{max}$ (µM) | $T_{max}$ | AUC |
|---|---|---|---|---|---|
| AME | Male | Blood | 3.82E-09 | 0.27338 | 2.45E-06 |
| | | Large Intestine Tissue | 5.35E-06 | 0.46836 | 6.82E-06 |
| | | Liver | 9.00E-08 | 0.17468 | 4.67E-08 |
| | | Kidney Tissue | 1.00E-08 | 0.1538 | 5.09E-09 |
| | Female | Blood | 4.41E-09 | 0.25816 | 2.59E-06 |
| | | Large Intestine Tissue | 4.84E-06 | 0.45168 | 6.19E-06 |
| | | Liver | 1.00E-07 | 0.16958 | 5.01E-08 |
| | | Kidney Tissue | 1.00E-08 | 0.13206 | 5.46E-09 |
| | | Uterus Tissue | 2.18E-09 | - | - |
| | Male child | Blood | 1.01E-08 | 0.18256 | 4.79E-06 |
| | | Large Intestine Tissue | 1.09E-06 | 0.35206 | 1.14E-06 |
| | | Liver | 2.30E-07 | 0.12012 | 9.80E-08 |
| | | Kidney Tissue | 2.00E-08 | 0.09812 | 2.10E-08 |
| | Male toddler | Blood | 1.21E-08 | 0.17992 | 4.63E-06 |
| | | Large Intestine Tissue | 1.95E-06 | 0.17616 | 1.04E-06 |
| | | Liver | 3.00E-07 | 0.11766 | 1.32E-07 |
| | | Kidney Tissue | 2.00E-08 | 0.1117 | 1.44E-08 |

**Table S6 *continued*:**

(C)

| Compound | Population | Compartment | $C_{max}$ (µM) | $T_{max}$ | AUC |
|---|---|---|---|---|---|
| AOH | Male | Blood | 1.70E-08 | 0.27068 | 9.94E-06 |
| | | Large Intestine Tissue | 5.12E-06 | 0.45172 | 6.40E-06 |
| | | Liver | 3.30E-07 | 0.17182 | 1.69E-07 |
| | | Kidney Tissue | 3.00E-08 | 0.17988 | 4.33E-08 |
| | Female | Blood | 1.96E-08 | 0.25652 | 1.05E-05 |
| | | Large Intestine Tissue | 4.64E-06 | 0.45276 | 5.81E-06 |
| | | Liver | 3.70E-07 | 0.17556 | 1.81E-07 |
| | | Adipose Tissue | 2.00E-08 | 0.78244 | 2.87E-07 |
| | | Kidney Tissue | 4.00E-08 | 0.23352 | 4.37E-08 |
| | | Uterus Tissue | 9.17E-09 | - | - |
| | Male child | Blood | 2.95E-08 | 0.1736 | 1.27E-05 |
| | | Large Intestine Tissue | 6.90E-07 | 0.33936 | 7.08E-07 |
| | | Liver | 5.60E-07 | 0.10978 | 2.32E-07 |
| | | Kidney Tissue | 5.00E-08 | 0.1151 | 6.94E-08 |
| | Male toddler | Blood | 4.19E-08 | 0.17284 | 1.45E-05 |
| | | Large Intestine Tissue | 1.44E-06 | 0.1666 | 7.62E-07 |
| | | Liver | 8.70E-07 | 0.11344 | 3.70E-07 |
| | | Kidney Tissue | 6.00E-08 | 0.15732 | 5.71E-08 |

**Table S6 *continued*:**

(C)

| Compound | Population | Compartment | $C_{max}$ (µM) | $T_{max}$ | AUC |
|---|---|---|---|---|---|
| AME | Male | Blood | 2.61E-09 | 0.26764 | 1.68E-06 |
| | | Large Intestine Tissue | 3.66E-06 | 0.46054 | 4.67E-06 |
| | | Liver | 6.00E-08 | 0.15396 | 3.16E-08 |
| | | Kidney Tissue | 1.06E-08 | 0.21874 | 2.37E-09 |
| | Female | Blood | 3.01E-09 | 0.25326 | 1.78E-06 |
| | | Large Intestine Tissue | 3.31E-06 | 0.44596 | 4.24E-06 |
| | | Liver | 7.00E-08 | 0.16958 | 3.40E-08 |
| | | Kidney Tissue | 7.90E-09 | 0.1777 | 3.10E-09 |
| | | Uterus Tissue | 1.50E-09 | - | - |
| | Male child | Blood | 6.17E-09 | 0.17918 | 2.92E-06 |
| | | Large Intestine Tissue | 6.60E-07 | 0.31928 | 6.96E-07 |
| | | Liver | 1.40E-07 | 0.10598 | 5.96E-08 |
| | | Kidney Tissue | 1.52E-08 | 0.06004 | 9.20E-09 |
| | Male toddler | Blood | 7.42E-09 | 0.17624 | 2.84E-06 |
| | | Large Intestine Tissue | 1.19E-06 | 0.1663 | 6.38E-07 |
| | | Liver | 1.80E-07 | 0.10104 | 8.06E-08 |
| | | Kidney Tissue | 1.41E-08 | 0.06592 | 7.07E-09 |

**Table S6 *continued*:**

(D)

| Compound | Population | Compartment | $C_{max}$ (µM) | $T_{max}$ | AUC |
|---|---|---|---|---|---|
| AOH | Male | Blood | 6.80E-08 | 0.28084 | 3.96E-05 |
| | | Large Intestine Tissue | 2.04E-05 | 0.46184 | 2.55E-05 |
| | | Liver | 1.33E-06 | 0.19312 | 6.76E-07 |
| | | Kidney Tissue | 1.30E-07 | 0.25586 | 2.73E-07 |
| | Female | Blood | 7.79E-08 | 0.25838 | 4.17E-05 |
| | | Large Intestine Tissue | 1.85E-05 | 0.45862 | 2.32E-05 |
| | | Liver | 1.47E-06 | 0.1848 | 7.24E-07 |
| | | Adipose Tissue | 8.00E-08 | 1.48236 | 1.12E-06 |
| | | Kidney Tissue | 1.40E-07 | 0.21538 | 2.84E-07 |
| | | Uterus Tissue | 3.66E-08 | - | - |
| | Male child | Blood | 1.51E-07 | 0.17492 | 6.49E-05 |
| | | Large Intestine Tissue | 3.52E-06 | 0.35802 | 3.62E-06 |
| | | Liver | 2.85E-06 | 0.11626 | 1.19E-06 |
| | | Kidney Tissue | 2.60E-07 | 0.14842 | 4.21E-07 |
| | Male toddler | Blood | 2.10E-07 | 0.176 | 7.23E-05 |
| | | Large Intestine Tissue | 7.21E-06 | 0.1808 | 3.81E-06 |
| | | Liver | 4.34E-06 | 0.12034 | 1.85E-06 |
| | | Kidney Tissue | 2.80E-07 | 0.15814 | 3.71E-07 |

**Table S6 *continued*:**

(D)

| Compound | Population | Compartment | $C_{max}$ (µM) | $T_{max}$ | AUC |
|---|---|---|---|---|---|
| AME | Male | Blood | 9.26E-09 | 0.29564 | 5.90E-06 |
| | | Large Intestine Tissue | 1.29E-05 | 0.46246 | 1.64E-05 |
| | | Liver | 2.10E-07 | 0.17548 | 1.13E-07 |
| | | Kidney Tissue | 2.00E-08 | 0.1837 | 1.71E-08 |
| | Female | Blood | 1.06E-08 | 0.26338 | 6.24E-06 |
| | | Large Intestine Tissue | 1.17E-05 | 0.47654 | 1.49E-05 |
| | | Liver | 2.40E-07 | 0.18818 | 1.22E-07 |
| | | Kidney Tissue | 2.00E-08 | 0.15446 | 1.80E-08 |
| | | Uterus Tissue | 5.28E-09 | - | - |
| | Male child | Blood | 2.41E-08 | 0.18068 | 1.14E-05 |
| | | Large Intestine Tissue | 2.60E-06 | 0.35762 | 2.73E-06 |
| | | Liver | 5.40E-07 | 0.11484 | 2.36E-07 |
| | | Kidney Tissue | 5.00E-08 | 0.138 | 6.89E-08 |
| | Male toddler | Blood | 2.78E-08 | 0.1859 | 1.06E-05 |
| | | Large Intestine Tissue | 4.46E-06 | 0.17862 | 2.39E-06 |
| | | Liver | 6.80E-07 | 0.11752 | 3.04E-07 |
| | | Kidney Tissue | 4.00E-08 | 0.11456 | 4.28E-08 |
