## Supplementary material for "Human internal exposures to alternariol and its monomethyl ether are predicted below thresholds of *in vitro* toxicity by physiologically based kinetic modeling": S2 Murine model for AOH.pdf

**Model S2:** Murine model for AOH

```
METHOD RK4

STARTTIME = 0
STOPTIME = 10
DT = 0.00002

{Type Equations Here.}
{Top model}
{Reservoirs}

d/dt (ASTAOH) = - transSTtoSI + oraldose
INIT ASTAOH = 0
LIMIT ASTAOH >= 0
d/dt (ASIAOH) = -transSItoSIG - transSItoLI  +  transSTtoSI
INIT ASIAOH = 0
LIMIT ASIAOH >= 0
d/dt (ALIAOH) =  +  transSItoLI - fecalex - transLItoL
INIT ALIAOH = 0
LIMIT ALIAOH >= 0
d/dt (AGSIAOH) =  + transSItoSIG  -venGS
INIT AGSIAOH = 0
LIMIT AGSIAOH >= 0
d/dt (FECAOH) = + fecalex
INIT FECAOH = 0
LIMIT FECAOH >= 0
d/dt (ALAOH) = - venL + artL + venP + venGS  + transLItoL - Phase1metab - AOHglucur
INIT ALAOH = 0
LIMIT ALAOH >= 0
d/dt (ABAOH) = + venL + venS + venR + venF + venK + venM  - artL - artP - artS - artR - artF - artK - artM
INIT ABAOH = 0
LIMIT ABAOH >= 0
d/dt (APAOH) = + artP - venP
INIT APAOH = 0
LIMIT APAOH >= 0
d/dt (ASAOH) = + artS - venS
INIT ASAOH = 0
LIMIT ASAOH >= 0
d/dt (ARAOH) = + artR - venR
INIT ARAOH = 0
LIMIT ARAOH >= 0
d/dt (AFAOH) = + artF - venF
INIT AFAOH = 0
LIMIT AFAOH >= 0
d/dt (AKAOH) = + artK - venK - urineAOH
INIT AKAOH = 0
LIMIT AKAOH >= 0
d/dt (AUCAOHUr) = + urineAOH
INIT AUCAOHUr = 0
d/dt (AMAOH) = + artM - venM
INIT AMAOH = 0
LIMIT AMAOH >= 0
d/dt (Start) = - oraldose
INIT Start = AODOSEAOH
LIMIT Start >=0
d/dt (exAOHmetab) = + Phase1metab
INIT exAOHmetab = 0
LIMIT exAOHmetab >= 0
d/dt (exAOHgluc) = + AOHglucur
INIT exAOHgluc = 0
LIMIT exAOHgluc >= 0
```

```
{Flows}

; arterial (art) and venous (ven) blood flows :

venL = QL*CVLAOH
artL = (QL-QP)*CBAOH
venP = QP*CPAOH
artP = QP*CBAOH
venS = QS*CVSAOH
artS = QS*CBAOH
venR = QR*CVRAOH
artR = QR*CBAOH
venF = QF*CVFAOH
artF = QF*CBAOH
venM = QM*CVMAOH
artM = QM*CBAOH
venK = QK*CVKAOH
artK = QK*CBAOH
venGS = QSI*CVGSIAOH
artGS = QSI*CBAOH

; excretion :

urineAOH = GFR*CVKAOH*60/1000  ;  (L/h)
fecalex = Kli*ALIAOH

; gastrointestinal tract :

transSTtoSI = kelST * ASTAOH              ; from stomach lumen to small intestine lumen
transSItoSIG = KaAOH*CSIAOH               ; from small intestine lumen to gut small intestine tissue
transSItoLI = ASIAOH*Ksi                  ; from small intestine lumen to large intestine lumen
transLItoL = KbAOH*CLIAOH                 ; from large intestine lumen to liver tissue

oraldose = PULSE(AODOSEAOH, 0, 24)

;Liver metabolism :
Phase1metab = VmaxLAOH * CVLAOH/(KmLAOH + CVLAOH)          ; MichaelisMenten
AOHglucur = VmaxLAOHgluc * CVLAOH/(KmLAOHgluc + CVLAOH)  ; MichaelisMenten

{Globals}
; AOH mouse model
;==================================================================
; Physiological parameters
;==================================================================
;bodyweight
BW = 0.037           ;Kg         ; avg. BW in validation study              ;EFSA in vivo study
;--------------------------------------------------------------------------------------------------------------
; relative tissue volumes
; fraction of BW
VLc = 0.0549                   ; fraction of liver tissue (Brown et al)
VPc = 0.0155                   ; fraction portal vein perfused tissue (stomach, spleen, pancreas) (Brown et al, Crispen et al)
VRc =0.0123                    ; fraction of rapidly perfused tis. (Q/V >>1, heart + lungs) (Brown et al)
VSc = 0.2726                   ; fraction of slowly perfused tissue (bone, skin) (Brown et al)
VFc = 0.0048                   ; fraction of fat tissue (adipose) (Brown et al)
VMc = 0.3840                   ; fraction of muscle tissue (Brown et al)
VKc = 0.0167                   ; fraction of kidneys (Brown et al)
VSIc = 0.0253                  ;fraction of small intestine tissue (Brown et al)
VLIc = 0.0109                  ;fraction of large intestine tissue (Brown et al)

;--------------------------------------------------------------------------------------------------------------
; calculated tissue volumes
VL = VLc*BW                ;L or Kg                     ; volume of liver tissue (calculated)
VP = VPc*BW                ;L or kg                     ; volume of portal vein perfused tissue
VR = VRc*BW                 ;L or Kg                    ; volume of rapidly perfused tissue (calculated)
```

```
VS = VSc*BW            ;L or Kg               ; volume of slowly perfused tissue (calculated)
VF = VFc*BW            ;L or Kg               ; volume of fat tissue (calculated)
VM = VMc*BW            ;L or Kg               ; volume of muscle tissue (calculated)
VK = VKc*BW            ;L or Kg               ; volume of kidney (calculated)
VB = 0.08*BW           ;L or Kg               ; volume of blood (Brown et al, calculated)
VSI = VSIc*BW          ;L or Kg               ; volume of small intestine tissue (calculated)
VLI = VLIc*BW          ;L or Kg               ; volume of large intestine tissue  (calculated)
VSIL = 0.002851        ;L or Kg               ; volume of small intestine lumen (Casteleyn et al, calculated)
VLIL = 0.001270        ;L or Kg               ; volume of large intestine lumen (Casteleyn et al, calculated)

;-------------------------------------------------------------------------------------------------------------------
;surface areas
areaSI = 0.3615    ;dm2                       ;surface area of SI lumen           ; (Casteleyn et al, calculated)
areaLI = 0.1234    ;dm2                       ;surface area of LI lumen    ; (Casteleyn et al, calculated)

;-------------------------------------------------------------------------------------------------------------------
;Blood flow rates,
QCc = 0.275*BW^0.75    ; (L/min), cardiac output (Brown et al)
QPc = 0.141            ; fraction of blood flow to portal vein (Brown et al)
QLc = 0.162            ; fraction of blood flow to liver in total (Brown et al)
QRc = 0.071            ; fraction of blood flow to rapidly perfused tissue (heart+lungs) (Brown et al)
QSc = 0.18             ; fraction of blood flow to slowly perfused tissue (skin+bones) (Brown et al)
QFc = 0.07             ; fraction of blood flow to fat (Brown et al)
QMc = 0.159            ; fraction of blood flow to muscle (Brown et al)
QKc = 0.091            ; fraction of blood flow to kidneys (Brown et al)
QSIc = 0.135           ; fraction of blood flow to small intestine (Stott et al)

QC = QCc*60            ; (L/h)                ; cardiac output (calculated)
QP = QPc*QC            ;L or Kg               ; blood flow to portal vein perfused tissue (calculated)
QL = QLc*QC            ;L or Kg               ; blood flow to liver total (calculated)
QR = QRc*QC            ;L or Kg               ; blood flow to rapidly perfused tissue (calculated)
QS = QSc*QC            ;L or Kg               ; blood flow to slowly perfused tissue (calculated)
QF = QFc*QC            ;L or Kg               ; blood flow to fat tissue (calculated)
QM = QMc*QC            ;L or Kg               ; blood flow to muscle tissue (calculated)
QK = QKc*QC            ;L or Kg               ; blood flow to kidney (calculated)
QSI=QSIc*QC            ;L or Kg               ; blood flow to small intestine tissue (calculated)

;-------------------------------------------------------------------------------------------------------------------
; glomerular filtration rate
GFR  = 0.28            ;mL/min                        ; GFR (Bivona et al)

;==================================================================
; Physicochemical parameters
;==================================================================
; molecular weights (g/mol)
MWAOH = 258.23 ; parent compound
; pKa1
pKa1AOH =7.63          ;Chemaxon playground (v1.6.2)
; pKa2
pKaAOH =8.44           ;Chemaxon playground (v1.6.2)
; logP
logPAOH = 3.18         ;Chemaxon playground (v1.6.2)
; logD (parent compound)
logDapical = 3.17      ;Chemaxon playground (v1.6.2)
logDbasal = 2.96       ;Chemaxon playground (v1.6.2)

; partition coefficients          ;calculated by qivivetools.wu.nl (Berezhkovskiy algorithm)
; Alternariol in main model
PLAOH = 2.2            ; liver/blood partition coefficient
PRAOH = 2.215          ; rapidly perfused tissue/blood partition coefficient
PSAOH = 2.621          ; slowly perfused tissue/blood partition coefficient
PFAOH = 7.9            ; fat/blood partition coefficient
PMAOH = 1.39           ; muscle/blood partition coefficient
```

```
PKAOH = 2.06                    ; kidney/blood partition coefficient
FUAOH = 0.06                    ; faction unbound plasma according to Lobell&Sivarajah
PSIAOH =3.07                    ; gut/blood partition coefficient
;--------------------------------------------------------------------------------------------------------------------
; absorption/transfer rates
Papp = 5.50E-06
logPappAOH = -5.09          ;log(cm/s)          ;log permeability calculated from Caco2 data (Burkhardt et al)
PeffAOH = (3600/10)*(10^(0.6836*LogPappAOH- 0.5579))          ;dm/h     ;scaled according to Sun et al.


KaAOH = PeffAOH*areaSI              ;L/h ;transfer rate of AOH from SI lumen to gut
KbAOH = PeffAOH*areaLI              ;L/h ;transfer rate of AOH from LI lumen to gut

; gastric emptying (kelst)

GEstc = 74                          ; (min) half time of meal in mice stomach (Schwarz et al)

GEst = GEstc /60                ; (h)
LNhalf= -0.69314718                 ; LN(1/2)
kelST =  -LNhalf/ GEst          ; (h-1) constant of gastric excretion                 ; from Ct=C0*e^(-kel*t)

tpassSI = 1.5                   ;h          ;passing time through small intestine (Woting et al)
Ksi = 1/tpassSI                 ;/h         ;SI to LI transfer constant
tpassLI = 8                     ;h          ;passing time through large intestine (Woting et al)
Kli = 1/tpassLI                 ;/h         ;LI to excretion transfer constant
;=====================================================================
; Kinetic parameters
;=====================================================================
;metabolism of rat liver
; scaling factors
VLS9 = 143          ;mg S9 protein/gram liver              ; liver S9 protein yield (Punt et al)
VLMP = 62           ;mg protein/gram liver                 ; liver microsomal protein yield (Smith et al)

;Phase 1 metabolism
KmLAOH = 78.7   ;µmol/L                 ;Michaelis constant of hepatic phase I metabolism (Borsos et al)
VmaxLAOHc = 0.00283                     ;µmol/mg/min   microsomal protein (Borsos et al)
VmaxLAOH = VmaxLAOHc*VLMP*VL*60*10^3          ;µmol/h

;Phase 2 glucuronidation
KmLAOHgluc = 8.5090       ;µmol/L               ;Michaelis constant of hepatic glucuronidation    (in vitro)
VmaxLAOHglucc =0.021437                 ;µmol/(mg*min) (in vitro)
VmaxLAOHgluc = VmaxLAOHglucc*VLS9*VL*60*10^3             ;µmol/h
;=====================================================================
; Run settings
;=====================================================================
;--------------------------------------------------------------------------------------------------------------------; oral dose
ODOSEAOH = 200000*BW ;µg                               ;oral dose alternariol, variable (EFSA 2014)
AODOSEAOH = ODOSEAOH/MWAOH          ;umol               ;oral dose AOH in umol
;=====================================================================
; Main model calculations/dynamics:
;=====================================================================
;--------------------------------------------------------------------------------------------------------------------
;ASIAOH: amount of AOH in small intestinal lumen compartment                ;µmol
CSIAOH=ASIAOH/VSIL                                                  ;umol/L
;--------------------------------------------------------------------------------------------------------------------;large intestine compartment
;ALIAOH: amount of AOH in large intestine lumen                             ;µmol
CLIAOH = ALIAOH/VLIL                                                ;umol/L
;--------------------------------------------------------------------------------------------------------------------
;gut small intestine tissue compartment
;AGSIAOH: amount of AOH in small intestine tissue                           ;µmol
CGSIAOH = AGSIAOH/VSI                                                      ;µmol/L
CVGSIAOH = CGSIAOH/PSIAOH
;--------------------------------------------------------------------------------------------------------------------
; liver compartment
```

```
; ALAOH: amount of AOH in liver ;µmol
CLAOH = ALAOH/VL                ;umol/L
CVLAOH = CLAOH/PLAOH                ;partition with liver tissue
;------------------------------------------------------------------------------------------------------------------
; portal vein perfused tissue compartment
; APAOH = amount of AOH in spleen tissue ;µmol
CPAOH = APAOH/VP                ;umol/L
;-----------------------------------------------------------------------------------------------------------------
; fat (adipose tissue) compartment
CFAOH = AFAOH/VF                ;umol/L
CVFAOH = CFAOH/PFAOH                ;partition with adipose tissue
;------------------------------------------------------------------------------------------------------------------
; muscle compartment
; AMAOH = amount of AOH in muscle tissue    ;µmol
CMAOH = AMAOH/VM                ;umol/L
CVMAOH = CMAOH/PMAOH                ;partition with muscle tissue
;------------------------------------------------------------------------------------------------------------------
; rapidly perfused tissue
CRAOH = ARAOH/VR                ;umol/L
CVRAOH = CRAOH/PRAOH                ;partition with muscle tissue
;------------------------------------------------------------------------------------------------------------------
; slowly perfused tissue
CSAOH = ASAOH/VS                ;umol/L
CVSAOH = CSAOH/PSAOH                ;partition with slowly perfused tissue
;------------------------------------------------------------------------------------------------------------------
; kidney
CKAOH = AKAOH/VK                ;umol/L
CVKAOH = CKAOH/PKAOH                ;partition with kidney tissue
;-----------------------------------------------------------------------------------------------------------------
; blood compartment
CBAOH=ABAOH/VB                ;umol/L
CVBAOH = CBAOH*FUAOH                ;free concentration in plasma
CBAOHug = CBAOH*MWAOH        ;µg/L = ng/mL                ;concentration in blood in ng/ml
;------------------------------------------------------------------------------------------------------------------
;=========================================================================
; Main model: mass balance calculation
;=========================================================================
Total = AODOSEAOH
Calculated = ASTAOH + ASIAOH  + ALIAOH + AGSIAOH  + ALAOH + ABAOH + AKAOH + ARAOH + ASAOH +
AFAOH + APAOH + AMAOH + AUCAOHUr + FECAOH +exAOHmetab + exAOHgluc

ERROR=((Total-Calculated)/(Total+1E-30))*100
MASSBAL=Total-Calculated + 1

{End Globals}
```
