## Supplementary material for "Human internal exposures to alternariol and its monomethyl ether are predicted below thresholds of *in vitro* toxicity by physiologically based kinetic modeling": S3 Male model for AOH.pdf

**Model S3:** Male model for AOH

```
METHOD RK4

STARTTIME = 0
STOPTIME = 10
DT = 0.00002

{Type Equations Here.}
{Top model}
 {Reservoirs}

d/dt (ASTAOH) = - transSTtoSI + oraldose
INIT ASTAOH = 0
LIMIT ASTAOH >= 0
d/dt (ASIAOH) = -transSItoSIG - transSItoLI  +  transSTtoSI
INIT ASIAOH = 0
LIMIT ASIAOH >= 0
d/dt (ALIAOH) =  +  transSItoLI - fecalex - transLItoL
INIT ALIAOH = 0
LIMIT ALIAOH >= 0
d/dt (AGSIAOH) =  + transSItoSIG  -venGS
INIT AGSIAOH = 0
LIMIT AGSIAOH >= 0
d/dt (FECAOH) = + fecalex
INIT FECAOH = 0
LIMIT FECAOH >= 0
d/dt (ALAOH) = - venL + artL + venP + venGS  + transLItoL - Phase1metab - AOHglucur
INIT ALAOH = 0
LIMIT ALAOH >= 0
d/dt (ABAOH) = + venL + venS + venR + venF + venK + venM  - artL - artP - artS - artR - artF - artK - artM
INIT ABAOH = 0
LIMIT ABAOH >= 0
d/dt (APAOH) = + artP - venP
INIT APAOH = 0
LIMIT APAOH >= 0
d/dt (ASAOH) = + artS - venS
INIT ASAOH = 0
LIMIT ASAOH >= 0
d/dt (ARAOH) = + artR - venR
INIT ARAOH = 0
LIMIT ARAOH >= 0
d/dt (AFAOH) = + artF - venF
INIT AFAOH = 0
LIMIT AFAOH >= 0
d/dt (AKAOH) = + artK - venK - urineAOH
INIT AKAOH = 0
LIMIT AKAOH >= 0
d/dt (AUCAOHUr) = + urineAOH
INIT AUCAOHUr = 0
d/dt (AMAOH) = + artM - venM
INIT AMAOH = 0
LIMIT AMAOH >= 0
d/dt (Start) = - oraldose
INIT Start = AODOSEAOH
LIMIT Start >=0
d/dt (exAOHmetab) = + Phase1metab
INIT exAOHmetab = 0
LIMIT exAOHmetab >= 0
d/dt (exAOHglucur) = + AOHglucur
INIT exAOHglucur = 0
LIMIT exAOHglucur >= 0

{Flows}
```

```
; arterial (art) and venous (ven) blood flows :

venL = QL*CVLAOH
artL = (QL-QP)*CBAOH
venP = QP*CPAOH
artP = QP*CBAOH
venS = QS*CVSAOH
artS = QS*CBAOH
venR = QR*CVRAOH
artR = QR*CBAOH
venF = QF*CVFAOH
artF = QF*CBAOH
venM = QM*CVMAOH
artM = QM*CBAOH
venK = QK*CVKAOH
artK = QK*CBAOH
venGS = QSI*CVGSIAOH
artGS = QSI*CBAOH

; excretion :

urineAOH = GFR*CVKAOH*60/1000  ;  (L/h)
fecalex = Kli*ALIAOH

; gastrointestinal tract :

transSTtoSI = kelST * ASTAOH            ; from stomach lumen to small intestine lumen
transSItoSIG = KaAOH*CSIAOH             ; from small intestine lumen to gut small intestine tissue
transSItoLI = ASIAOH*Ksi                ; from small intestine lumen to large intestine lumen
transLItoL = KbAOH*CLIAOH               ; from large intestine lumen to liver tissue

oraldose = PULSE(AODOSEAOH, 0, 24)

;Liver metabolism :
Phase1metab = VmaxLAOH * CVLAOH/(KmLAOH + CVLAOH)          ; MichaelisMenten
AOHglucur = VmaxLAOHgluc * CVLAOH/(KmLAOHgluc + CVLAOH)  ; MichaelisMenten

{Globals}
; AOH male model
;=====================================================================
; Physiological parameters ;=====================================================================
;bodyweight
BW = 73 ;Kg         ; avg. BW in validation study (ICRP)
;---------------------------------------------------------------------------------------------------------------
; relative tissue volumes
; fraction of BW
VLc = 0.0319            ; fraction of liver tissue (ICRP)
VPc = 0.0063            ; fraction portal vein perfused tissue (stomach, spleen, pancreas) tissue (ICRP)
VRc =0.0212             ; fraction of rapidly perfused tis. (Q/V >>1, heart + lungs) tissue (ICRP)
VSc = 0.1963            ; fraction of slowly perfused tissue (bone, skin) tissue (ICRP)
VFc = 0.2036            ; fraction of fat tissue (adipose) tissue (ICRP)
VMc = 0.4074            ; fraction of muscle tissue tissue (ICRP)
VKc = 0.0057            ; fraction of kidneys tissue (ICRP)
VSIc = 0.0117           ; fraction of small intestine tissue (ICRP)
VLIc = 0.0067           ; fraction of large intestine tissue (ICRP)
VBc = 0.0726            ; fraction of blood (ICRP)

;---------------------------------------------------------------------------------------------------------------
; calculated tissue volumes
VL = VLc*BW             ;L or Kg            ; volume of liver tissue (calculated)
VP = VPc*BW             ;L or kg            ; volume of portal vein perfused tissue
VR = VRc*BW             ;L or Kg            ; volume of rapidly perfused tissue (calculated)
VS = VSc*BW             ;L or Kg            ; volume of slowly perfused tissue (calculated)
```

```
VF = VFc*BW              ;L or Kg                 ; volume of fat tissue (calculated)
VM = VMc*BW              ;L or Kg                 ; volume of muscle tissue (calculated)
VK = VKc*BW              ;L or Kg                 ; volume of kidney (calculated)
VB = VBc*BW              ;L or Kg                 ; volume of blood (Brown et al, calculated)
VSI = VSIc*BW            ;L or Kg                 ; volume of small intestine tissue (calculated)
VLI = VLIc*BW            ;L or Kg                 ; volume of large intestine tissue  (calculated)
VSIL = 4.45              ;L or Kg                 ; volume of small intestine lumen (ICPR, calculated)
VLIL = 2.38              ;L or Kg                 ; volume of large intestine lumen (ICRP, calculated)
;------------------------------------------------------------------------------------------------------------------
;surface areas

areaSI = 2917.5 ;dm2     ;surface area of SI lumen (Helander et al)
areaLI = 107.8;dm2                 ;surface area of LI lumen (Helander et al)
;------------------------------------------------------------------------------------------------------------------
;Blood flow rates,
QCc = 6.5                ; (L/min), cardiac output (ICRP)
QPc = 0.05               ; fraction of blood flow to P (spleen, stom., pancreas) (ICRP)
QLc = 0.065              ; fraction of blood flow to liver in total (ICRP)
QRc = 1.04               ; fraction of blood flow to rapidly perfused tissue (heart+lungs)       (ICRP)
QSc = 0.1                ; fraction of blood flow to slowly perfused tissue (skin+Bones)         (ICRP)
QFc = 0.05               ; fraction of blood flow to fat  (ICRP)
QMc =0.17                ; fraction of blood flow to muscle (ICRP)
QKc =0.19                ; fraction of blood flow to kidneys (ICRP)
QSIc =0.1                ; fraction of blood flow to small intestine (ICRP)

QC = QCc*60              ; (L/h)          ; cardiac output (calculated)
QP = QPc*QC              ;L or Kg  ; blood flow to portal vein perfused tissue (calculated)
QL = QLc*QC              ;L or Kg  ; blood flow to liver total (calculated)
QR = QRc*QC              ;L or Kg  ; blood flow to rapidly perfused tissue (calculated)
QS = QSc*QC              ;L or Kg  ; blood flow to slowly perfused tissue (calculated)
QF = QFc*QC              ;L or Kg  ; blood flow to fat tissue (calculated)
QM = QMc*QC              ;L or Kg  ; blood flow to muscle tissue (calculated)
QK = QKc*QC              ;L or Kg  ; blood flow to kidney (calculated)
QSI=QSIc*QC              ;L or Kg  ; blood flow to small intestine tissue (calculated)
;------------------------------------------------------------------------------------------------------------------
; glomerular filtration rate
GFR = 125        ;mL/min          ; GFR scaled to body weight (Levey et al.)
;=================================================================
; Physicochemical parameters
;=================================================================
; molecular weights (g/mol)
MWAOH = 258.23 ; parent compound
; pKa1
pKa1AOH =7.63 ; parent compound Chemaxon playground (v1.6.2)
; pKa2
pKaAOH =8.44 ; parent compound Chemaxon playground (v1.6.2)
; logP
logPAOH = 3.18 ; parent compound Chemaxon playground (v1.6.2)
; logD (parent compound)
logDapical = 3.17 ;Chemaxon playground (v1.6.2)
logDbasal = 2.96 ; Chemaxon playground (v1.6.2)

; partition coefficients             ;calculated by qivivetools.wu.nl (Berezhkovskiy algorithm)
; Alternariol in main model
PLAOH = 3.21             ; liver/blood partition coefficient
PRAOH = 0.893            ; rapidly perfused tissue/blood partition coefficient
PSAOH = 4.601            ; slowly perfused tissue/blood partition coefficient
PFAOH = 7.3              ; fat/blood partition coefficient
PMAOH = 2.09             ; muscle/blood partition coefficient
PKAOH = 2.07             ; kidney/blood partition coefficient
FUAOH = 0.06             ; faction unbound plasma according to Lobell&Sivarajah
PSIAOH =3.97             ; gut/blood partition coefficient
;------------------------------------------------------------------------------------------------------------------
; absorption/transfer rates
```

```
logPappAOH = -5.09                    ;log(cm/s)          ;log permeability calculated from Caco2 data (Burkhardt et al)
PeffAOH = (3600/10)*(10^(0.6836*LogPappAOH- 0.5579))          ;dm/h     ;scaled according toSun et al.


KaAOH = PeffAOH*areaSI                ;L/h ;transfer rate of AOH from SI lumen to gut
KbAOH = PeffAOH*areaLI                ;L/h ;transfer rate of AOH from LI lumen to gut

; gastric emptying (kelst)
GEstc = 11.8                          ; (min) half time of meal in stomach
GEst = GEstc /60              ; (h)
LNhalf= -0.69314718                   ; LN(1/2)
kelST =  -LNhalf/ GEst        ; (h-1) constant of gastric excretion               ; from Ct=C0*e^(-kel*t)

tpassSI = 4                   ;h          ;passing time through small intestine (ICRP)
Ksi = 1/tpassSI               ;/h                   ;SI to LI transfer constant
tpassLI = 36                  ;h                    ;passing time through large intestine (ICRP)
Kli = 1/tpassLI               ;/h                   ;LI to excretion transfer constant
;===========================================================
; Kinetic parameters
;===========================================================
;metabolism of rat liver
; scaling factors
VLS9 = 107.3        ;mg S9 protein/gram liver               ; liver S9 protein yield (Wang et al. 2019)
VLMP = 32           ;mg protein/gram liver                  ; liver microsomal protein yield (Barter et al. 2007)

;Phase 1 metabolism
KmLAOH = 131                  ;µmol/L  ;Michaelis constant of hepatic phase I metabolism          (Borsos et al)
VmaxLAOHc = 0.00337                   ;µmol/mg/min   microsomal protein (Borsos et al)
VmaxLAOH = VmaxLAOHc*VLMP*VL*60*10^3          ;µmol/h

;Phase 2 glucuronidation
KmLAOHgluc = 2.7                                ;µmol/L            ;Michaelis constant of hepatic glucuronidation (in vitro)
VmaxLAOHglucc =0.0069                 ;µmol/(mg*min) (in vitro)
VmaxLAOHgluc = VmaxLAOHglucc*VLS9*VL*60*10^3               ;µmol/h
;===========================================================
; Run settings
;============================================================
;-----------------------------------------------------------------------------------------------------------
; oral dose
ODOSEAOH = 0.0022*BW  ;µg                             ;oral dose alternariol, variable
AODOSEAOH = ODOSEAOH/MWAOH           ;umol                  ;oral dose AOH in umol
;============================================================
; Main model calculations/dynamics:
;============================================================
;-----------------------------------------------------------------------------------------------------------
;ASIAOH: amount of AOH in small intestinal lumen compartment                ;µmol
CSIAOH=ASIAOH/VSIL                                         ;umol/L
;-----------------------------------------------------------------------------------------------------------
;large intestine compartment
;ALIAOH: amount of AOH in large intestine lumen                             ;µmol
CLIAOH = ALIAOH/VLIL                                       ;umol/L

;-----------------------------------------------------------------------------------------------------------;gut small intestine tissue

compartment

;AGSIAOH: amount of AOH in small intestine tissue                           ;µmol

CGSIAOH = AGSIAOH/VSI                                                      ;µmol/L

CVGSIAOH = CGSIAOH/PSIAOH
;-----------------------------------------------------------------------------------------------------------
; liver compartment
; ALAOH: amount of AOH in liver ;µmol
CLAOH = ALAOH/VL                      ;umol/L
```

```
CVLAOH = CLAOH/PLAOH            ;partition with liver tissue
;---------------------------------------------------------------------------------------------------------------
; portal vein perfused tissue compartment
; APAOH = amount of AOH in spleen tissue ;µmol
CPAOH = APAOH/VP                ;umol/L
;---------------------------------------------------------------------------------------------------------------
; fat (adipose tissue) compartment
CFAOH = AFAOH/VF                ;umol/L
CVFAOH = CFAOH/PFAOH            ;partition with adipose tissue
;---------------------------------------------------------------------------------------------------------------
; muscle compartment
; AMAOH = amount of AOH in muscle tissue     ;µmol
CMAOH = AMAOH/VM                ;umol/L
CVMAOH = CMAOH/PMAOH            ;partition with muscle tissue
;---------------------------------------------------------------------------------------------------------------
; rapidly perfused tissue
CRAOH = ARAOH/VR                ;umol/L
CVRAOH = CRAOH/PRAOH            ;partition with muscle tissue
;---------------------------------------------------------------------------------------------------------------
; slowly perfused tissue
CSAOH = ASAOH/VS                ;umol/L
CVSAOH = CSAOH/PSAOH            ;partition with slowly perfused tissue
;---------------------------------------------------------------------------------------------------------------
; kidney
CKAOH = AKAOH/VK                ;umol/L
CVKAOH = CKAOH/PKAOH            ;partition with kidney tissue
;---------------------------------------------------------------------------------------------------------------
; blood compartment
CBAOH=ABAOH/VB                  ;umol/L
CVBAOH = CBAOH*FUAOH            ;free concentration in plasma
CBAOHug = CBAOH*MWAOH           ;µg/L = ng/mL            ;concentration in blood in ng/ml
;---------------------------------------------------------------------------------------------------------------
;=====================================================================
; Main model: mass balance calculation
;=====================================================================
Total = AODOSEAOH
Calculated = ASTAOH + ASIAOH  + ALIAOH + AGSIAOH  + ALAOH + ABAOH + AKAOH + ARAOH + ASAOH +
AFAOH + APAOH + AMAOH + AUCAOHUr + FECAOH +exAOHmetab +exAOHglucur

ERROR=((Total-Calculated)/(Total+1E-30))*100
MASSBAL=Total-Calculated + 1

{End Globals}
```
