## Supplementary material for "Human internal exposures to alternariol and its monomethyl ether are predicted below thresholds of *in vitro* toxicity by physiologically based kinetic modeling": S5 Male child model for AOH.pdf

**Model S5:** Male child model for AOH

METHOD RK4

STARTTIME = 0
STOPTIME = 10
DT = 0.00002

{Type Equations Here.}
{Top model}
{Reservoirs}

d/dt (ASTAOH) = - transSTtoSI + oraldose
INIT ASTAOH = 0
LIMIT ASTAOH >= 0
d/dt (ASIAOH) = -transSItoSIG - transSItoLI + transSTtoSI
INIT ASIAOH = 0
LIMIT ASIAOH >= 0
d/dt (ALIAOH) = + transSItoLI - fecalex - transLItoL
INIT ALIAOH = 0
LIMIT ALIAOH >= 0
d/dt (AGSIAOH) = + transSItoSIG -venGS
INIT AGSIAOH = 0
LIMIT AGSIAOH >= 0
d/dt (FECAOH) = + fecalex
INIT FECAOH = 0
LIMIT FECAOH >= 0
d/dt (ALAOH) = - venL + artL + venP + venGS + transLItoL - Phase1metab - AOHglucur
INIT ALAOH = 0
LIMIT ALAOH >= 0
d/dt (ABAOH) = + venL + venS + venR + venF + venK + venM - artL - artP - artS - artR - artF - artK - artM
INIT ABAOH = 0
LIMIT ABAOH >= 0
d/dt (APAOH) = + artP - venP
INIT APAOH = 0
LIMIT APAOH >= 0
d/dt (ASAOH) = + artS - venS
INIT ASAOH = 0
LIMIT ASAOH >= 0
d/dt (ARAOH) = + artR - venR
INIT ARAOH = 0
LIMIT ARAOH >= 0
d/dt (AFAOH) = + artF - venF
INIT AFAOH = 0
LIMIT AFAOH >= 0
d/dt (AKAOH) = + artK - venK - urineAOH
INIT AKAOH = 0
LIMIT AKAOH >= 0
d/dt (AUCAOHUr) = + urineAOH
INIT AUCAOHUr = 0
d/dt (AMAOH) = + artM - venM
INIT AMAOH = 0
LIMIT AMAOH >= 0
d/dt (Start) = - oraldose
INIT Start = AODOSEAOH
LIMIT Start >=0
d/dt (exAOHmetab) = + Phase1metab
INIT exAOHmetab = 0
LIMIT exAOHmetab >= 0
d/dt (exAOHglucur) = + AOHglucur
INIT exAOHglucur = 0
LIMIT exAOHglucur >= 0

{Flows}

```
; arterial (art) and venous (ven) blood flows :

venL = QL*CVLAOH
artL = (QL-QP)*CBAOH
venP = QP*CPAOH
artP = QP*CBAOH
venS = QS*CVSAOH
artS = QS*CBAOH
venR = QR*CVRAOH
artR = QR*CBAOH
venF = QF*CVFAOH
artF = QF*CBAOH
venM = QM*CVMAOH
artM = QM*CBAOH
venK = QK*CVKAOH
artK = QK*CBAOH
venGS = QSI*CVGSIAOH
artGS = QSI*CBAOH

; excretion :

urineAOH = GFR*CVKAOH*60/1000  ;  (L/h)
fecalex = Kli*ALIAOH

; gastrointestinal tract :

transSTtoSI = kelST * ASTAOH          ; from stomach lumen to small intestine lumen
transSItoSIG = KaAOH*CSIAOH           ; from small intestine lumen to gut small intestine tissue
transSItoLI = ASIAOH*Ksi              ; from small intestine lumen to large intestine lumen
transLItoL = KbAOH*CLIAOH             ; from large intestine lumen to liver tissue

oraldose = PULSE(AODOSEAOH, 0, 24)

;Liver metabolism :
Phase1metab = VmaxLAOH * CVLAOH/(KmLAOH + CVLAOH)          ; MichaelisMenten
AOHglucur = VmaxLAOHgluc * CVLAOH/(KmLAOHgluc + CVLAOH)  ; MichaelisMenten

{Globals}
; AOH male child model
;==================================================================
; Physiological parameters ;==================================================================
;bodyweight
BW = 19 ;Kg          ; avg. BW in validation study (ICRP)
;-----------------------------------------------------------------------------------------------------------------
; relative tissue volumes
; fraction of BW
VLc = 0.0374            ; fraction of liver tissue (ICRP)
VPc = 0.0068            ; fraction portal vein perfused tissue (stomach, spleen, pancreas) tissue (ICRP)
VRc = 0.0210            ; fraction of rapidly perfused tis. (Q/V >>1, heart + lungs) tissue (ICRP)
VSc = 0.1653            ; fraction of slowly perfused tissue (bone, skin) tissue (ICRP)
VFc = 0.1932            ; fraction of fat tissue (adipose) tissue (ICRP)
VMc = 0.3051            ; fraction of muscle tissue tissue (ICRP)
VKc = 0.0073            ; fraction of kidneys tissue (ICRP)
VSIc = 0.0144           ; fraction of small intestine tissue (ICRP)
VLIc = 0.0079           ; fraction of large intestine tissue (ICRP)
VBc = 0.0737            ; fraction of blood (ICRP)

;-----------------------------------------------------------------------------------------------------------------
; calculated tissue volumes
VL = VLc*BW      ;L or Kg         ; volume of liver tissue (calculated)
VP = VPc*BW      ;L or kg         ; volume of portal vein perfused tissue
VR = VRc*BW      ;L or Kg         ; volume of rapidly perfused tissue (calculated)
VS = VSc*BW      ;L or Kg         ; volume of slowly perfused tissue (calculated)
```

```
VF = VFc*BW     ;L or Kg        ; volume of fat tissue (calculated)
VM = VMc*BW     ;L or Kg        ; volume of muscle tissue (calculated)
VK = VKc*BW     ;L or Kg        ; volume of kidney (calculated)
VB = VBc*BW     ;L or Kg        ; volume of blood (Brown et al, calculated)
VSI = VSIc*BW   ;L or Kg        ; volume of small intestine tissue (calculated)
VLI = VLIc*BW   ;L or Kg        ; volume of large intestine tissue  (calculated)
VSIL = 0.48     ;L or Kg        ; volume of small intestine lumen (ICPR, calculated)
VLIL = 1.19     ;L or Kg        ; volume of large intestine lumen (ICRP, calculated)
;---------------------------------------------------------------------------------------------------------------
;surface areas

areaSI = 1771.312;dm2       ;surface area of SI lumen (Helander et al)
areaLI = 73.51;dm2                  ;surface area of LI lumen (Helander et al)
;---------------------------------------------------------------------------------------------------------------
;Blood flow rates,
QCc = 3.4       ; (L/min), cardiac output (ICRP)
QPc = 0.05      ; fraction of blood flow to P (spleen, stom., pancreas) (ICRP)
QLc = 0.065     ; fraction of blood flow to liver in total (ICRP)
QRc = 1.04      ; fraction of blood flow to rapidly perfused tissue (heart+lungs)       (ICRP)
QSc = 0.1       ; fraction of blood flow to slowly perfused tissue (skin+Bones)         (ICRP)
QFc = 0.05      ; fraction of blood flow to fat  (ICRP)
QMc =0.17       ; fraction of blood flow to muscle (ICRP)
QKc =0.19       ; fraction of blood flow to kidneys (ICRP)
QSIc =0.1       ; fraction of blood flow to small intestine (ICRP)
QLIc=0.04       ; fraction of blood flow to large intestine (ICRP)

QC = QCc*60             ; (L/h)         ; cardiac output (calculated)
QP = QPc*QC             ;L or Kg        ; blood flow to portal vein perfused tissue (calculated)
QL = QLc*QC             ;L or Kg        ; blood flow to liver total (calculated)
QR = QRc*QC             ;L or Kg        ; blood flow to rapidly perfused tissue (calculated)
QS = QSc*QC             ;L or Kg        ; blood flow to slowly perfused tissue (calculated)
QF = QFc*QC             ;L or Kg        ; blood flow to fat tissue (calculated)
QM = QMc*QC             ;L or Kg        ; blood flow to muscle tissue (calculated)
QK = QKc*QC             ;L or Kg        ; blood flow to kidney (calculated)
QSI=QSIc*QC             ;L or Kg        ; blood flow to small intestine tissue (calculated)
;---------------------------------------------------------------------------------------------------------------
; glomerular filtration rate
GFR = 125       ;mL/min         ; GFR scaled to body weight (Levey et al.)
;=================================================================
; Physicochemical parameters
;=================================================================
; molecular weights (g/mol)
MWAOH = 258.23 ; parent compound
; pKa1
pKa1AOH =7.63 ; parent compound Chemaxon playground (v1.6.2)
; pKa2
pKaAOH =8.44 ; parent compound Chemaxon playground (v1.6.2)
; logP
logPAOH = 3.18 ; parent compound Chemaxon playground (v1.6.2)
; logD (parent compound)
logDapical = 3.17 ;Chemaxon playground (v1.6.2)
logDbasal = 2.96 ; Chemaxon playground (v1.6.2)

; partition coefficients                ;calculated by qivivetools.wu.nl (Berezhkovskiy algorithm)
; Alternariol in main model
PLAOH = 3.21            ; liver/blood partition coefficient
PRAOH = 0.898           ; rapidly perfused tissue/blood partition coefficient
PSAOH = 4.733           ; slowly perfused tissue/blood partition coefficient
PFAOH = 7.3             ; fat/blood partition coefficient
PMAOH = 2.09            ; muscle/blood partition coefficient
PKAOH = 2.07            ; kidney/blood partition coefficient
FUAOH = 0.06            ; faction unbound plasma according to Lobell&Sivarajah
PSIAOH =3.97            ; gut/blood partition coefficient
;---------------------------------------------------------------------------------------------------------------
```

```
; absorption/transfer rates
logPappAOH = -5.09                    ;log(cm/s)          ;log permeability calculated from Caco2 data (Burkhardt et al)
PeffAOH = (3600/10)*(10^(0.6836*LogPappAOH- 0.5579))          ;dm/h     ;scaled according toSun et al.


KaAOH = PeffAOH*areaSI                ;L/h ;transfer rate of AOH from SI lumen to gut
KbAOH = PeffAOH*areaLI                ;L/h ;transfer rate of AOH from LI lumen to gut

; gastric emptying (kelst)
GEstc = 11.8                          ; (min) half time of meal in stomach
GEst = GEstc /60               ; (h)
LNhalf= -0.69314718                   ; LN(1/2)
kelST =  -LNhalf/ GEst         ; (h-1) constant of gastric excretion                 ; from Ct=C0*e^(-kel*t)

tpassSI = 4                    ;h            ;passing time through small intestine (ICRP)
Ksi = 1/tpassSI                ;/h                  ;SI to LI transfer constant
tpassLI = 34                   ;h                   ;passing time through large intestine (ICRP)
Kli = 1/tpassLI                ;/h                  ;LI to excretion transfer constant
;==============================================================
; Kinetic parameters
;==============================================================
;metabolism of rat liver
; scaling factors
VLS9 = 107.3        ;mg S9 protein/gram liver              ; liver S9 protein yield (Wang et al. 2019)
VLMP = 32           ;mg protein/gram liver                 ; liver microsomal protein yield (Barter et al. 2007)

;Phase 1 metabolism
KmLAOH = 131                   ;µmol/L   ;Michaelis constant of hepatic phase I metabolism          (Borsos et al)
VmaxLAOHc = 0.00337                   ;µmol/mg/min   microsomal protein (Borsos et al)
VmaxLAOH = VmaxLAOHc*VLMP*VL*60*10^3          ;µmol/h

;Phase 2 glucuronidation
KmLAOHgluc = 2.7                      ;µmol/L             ;Michaelis constant of hepatic glucuronidation (in vitro)
VmaxLAOHglucc =0.0069                 ;µmol/(mg*min) (in vitro)
VmaxLAOHgluc = VmaxLAOHglucc*VLS9*VL*60*10^3           ;µmol/h
;==============================================================
; Run settings
;==============================================================
;------------------------------------------------------------------------------------------------------------
; oral dose
ODOSEAOH = 0.0031*BW  ;µg                          ;oral dose alternariol, variable
AODOSEAOH = ODOSEAOH/MWAOH          ;umol                ;oral dose AOH in umol
;==============================================================
; Main model calculations/dynamics:
;==============================================================
;------------------------------------------------------------------------------------------------------------
;ASIAOH: amount of AOH in small intestinal lumen compartment   ;µmol
CSIAOH=ASIAOH/VSIL                                              ;umol/L
;------------------------------------------------------------------------------------------------------------
;large intestine compartment
;ALIAOH: amount of AOH in large intestine lumen                 ;µmol
CLIAOH = ALIAOH/VLIL                                            ;umol/L

;-----------------------------------------------------------------------------------------------------------;gut small intestine tissue

compartment

;AGSIAOH: amount of AOH in small intestine tissue               ;µmol

CGSIAOH = AGSIAOH/VSI                                           ;µmol/L

CVGSIAOH = CGSIAOH/PSIAOH
;------------------------------------------------------------------------------------------------------------
; liver compartment
; ALAOH: amount of AOH in liver ;µmol
```
