## Supplementary material for "Human internal exposures to alternariol and its monomethyl ether are predicted below thresholds of *in vitro* toxicity by physiologically based kinetic modeling": S7 Male model for AME.pdf

**Model S7:** Male model for AME

METHOD RK4

STARTTIME = 0
STOPTIME = 10
DT = 0.00002

{Type Equations Here.}
{Top model}
{Reservoirs}

d/dt (ASTAME) = - transSTtoSI + oraldose
INIT ASTAME = 0
LIMIT ASTAME >= 0
d/dt (ASIAME) = -transSItoSIG - transSItoLI + transSTtoSI
INIT ASIAME = 0
LIMIT ASIAME >= 0
d/dt (ALIAME) = + transSItoLI - fecalex - transLItoL
INIT ALIAME = 0
LIMIT ALIAME >= 0
d/dt (AGSIAME) = + transSItoSIG -venGS
INIT AGSIAME = 0
LIMIT AGSIAME >= 0
d/dt (FECAME) = + fecalex
INIT FECAME = 0
LIMIT FECAME >= 0
d/dt (ALAME) = - venL + artL + venP + venGS + transLItoL - Phase1metab - AMEglucur
INIT ALAME = 0
LIMIT ALAME >= 0
d/dt (ABAME) = + venL + venS + venR + venF + venK + venM - artL - artP - artS - artR -artF - artK - artM
INIT ABAME = 0
LIMIT ABAME >= 0
d/dt (APAME) = + artP - venP
INIT APAME = 0
LIMIT APAME >= 0
d/dt (ASAME) = + artS - venS
INIT ASAME = 0
LIMIT ASAME >= 0
d/dt (ARAME) = + artR - venR
INIT ARAME = 0
LIMIT ARAME >= 0
d/dt (AFAME) = + artF - venF
INIT AFAME = 0
LIMIT AFAME >= 0
d/dt (AKAME) = + artK - venK - urineAME
INIT AKAME = 0
LIMIT AKAME >= 0
d/dt (AUCAMEUr) = + urineAME
INIT AUCAMEUr = 0
d/dt (AMAME) = + artM - venM
INIT AMAME = 0
LIMIT AMAME >= 0
d/dt (Start) = - oraldose
INIT Start = AODOSEAME
LIMIT Start >=0
d/dt (exAMEmetab) = + Phase1metab
INIT exAMEmetab = 0
LIMIT exAMEmetab >= 0
d/dt (exAMEglucur) = + AMEglucur
INIT exAMEglucur = 0
LIMIT exAMEglucur >= 0

{Flows}

```
 ; arterial (art) and venous (ven) blood flows :
venL = QL*CVLAME
artL = (QL-QP)*CBAME
venP = QP*CPAME
artP = QP*CBAME
venS = QS*CVSAME
artS = QS*CBAME
venR = QR*CVRAME
artR = QR*CBAME
venF = QF*CVFAME
artF = QF*CBAME
venM = QM*CVMAME
artM = QM*CBAME
venK = QK*CVKAME
artK = QK*CBAME
venGS = QSI*CVGSIAME
artGS = QSI*CBAME

 ; excretion :
urineAME = GFR*CVKAME*60/1000  ;  (L/h)
fecalex = Kli*ALIAME

; gastrointestinal tract :
transSTtoSI = kelST * ASTAME              ; from stomach lumen to small intestine lumen
transSItoSIG = KaAME*CSIAME               ; from small intestine lumen to gut small intestine tissue
transSItoLI = ASIAME*Ksi                  ; from small intestine lumen to large intestine lumen
transLItoL = KbAME*CLIAME                 ; from large intestine lumen to liver tissue

oraldose = PULSE(AODOSEAME, 0, 24)

;Liver metabolism :
Phase1metab = VmaxLAME * CVLAME/(KmLAME + CVLAME)          ; MichaelisMenten
AMEglucur = VmaxLAMEgluc * CVLAME/(KmLAMEgluc + CVLAME)  ; MichaelisMenten

{Globals}

;AME male model
;=================================================================
; Physiological parameters
;=================================================================
;bodyweight
BW = 73 ;Kg         ; avg. BW in validation study (ICRP)
;-----------------------------------------------------------------------------------------------------------------
; relative tissue volumes
; fraction of BW
VLc = 0.0319             ; fraction of liver tissue (ICRP)
VPc = 0.0063             ; fraction portal vein perfused tissue (stomach, spleen, pancreas) tissue (ICRP)
VRc =0.0212              ; fraction of rapidly perfused tis. (Q/V >>1, heart + lungs) tissue (ICRP)
VSc = 0.1963             ; fraction of slowly perfused tissue (bone, skin) tissue (ICRP)
VFc = 0.2036             ; fraction of fat tissue (adipose) tissue (ICRP)
VMc = 0.4074             ; fraction of muscle tissue tissue (ICRP)
VKc = 0.0057             ; fraction of kidneys tissue (ICRP)
VSIc = 0.0117            ; fraction of small intestine tissue (ICRP)
VLIc = 0.0067            ; fraction of large intestine tissue (ICRP)
VBc = 0.0726             ; fraction of blood (ICRP)


;-----------------------------------------------------------------------------------------------------------------
; calculated tissue volumes
VL = VLc*BW              ;L or Kg             ; volume of liver tissue (calculated)
VP = VPc*BW              ;L or kg             ; volume of portal vein perfused tissue
VR = VRc*BW              ;L or Kg             ; volume of rapidly perfused tissue (calculated)
VS = VSc*BW              ;L or Kg             ; volume of slowly perfused tissue (calculated)
VF = VFc*BW              ;L or Kg             ; volume of fat tissue (calculated)
```

VM = VMc*BW ;L or Kg ; volume of muscle tissue (calculated)
VK = VKc*BW ;L or Kg ; volume of kidney (calculated)
VB = VBc*BW ;L or Kg ; volume of blood (Brown et al, calculated)
VSI = VSIc*BW ;L or Kg ; volume of small intestine tissue (calculated)
VLI = VLIc*BW ;L or Kg ; volume of large intestine tissue (calculated)
VSIL = 4.45 ;L or Kg ; volume of small intestine lumen (ICPR, calculated)
VLIL = 2.38 ;L or Kg ; volume of large intestine lumen (ICPR, calculated);-----------------
---------------------------------------------------------------------------------------------------
;surface areas
areaSI = 2917.5 ;dm2 ;surface area of SI lumen (Helander et al.)
areaLI = 107.8 ;dm2 ;surface area of LI lumen (Helander et al.)
;-------------------------------------------------------------------------------------------------------------------
;Blood flow rates,
QCc = 6.5 ; (L/min), cardiac output (ICRP)
QPc = 0.05 ; fraction of blood flow to P (spleen, stom., pancreas) (ICRP)
QLc = 0.065 ; fraction of blood flow to liver in total (ICRP)
QRc = 1.04 ; fraction of blood flow to rapidly perfused tissue (heart+lungs) (ICRP)
QSc = 0.1 ; fraction of blood flow to slowly perfused tissue (skin+Bones) (ICRP)
QFc = 0.05 ; fraction of blood flow to fat (ICRP)
QMc =0.17 ; fraction of blood flow to muscle (ICRP)
QKc =0.19 ; fraction of blood flow to kidneys (ICRP)
QSIc =0.1 ; fraction of blood flow to small intestine (ICRP)
QLIc=0.04 ; fraction of blood flow to large intestine (ICRP)

QC = QCc*60 ; (L/h) ; cardiac output (calculated)
QP = QPc*QC ;L or Kg ; blood flow to portal vein perfused tissue (calculated)
QL = QLc*QC ;L or Kg ; blood flow to liver total (calculated)
QR = QRc*QC ;L or Kg ; blood flow to rapidly perfused tissue (calculated)
QS = QSc*QC ;L or Kg ; blood flow to slowly perfused tissue (calculated)
QF = QFc*QC ;L or Kg ; blood flow to fat tissue (calculated)
QM = QMc*QC ;L or Kg ; blood flow to muscle tissue (calculated)
QK = QKc*QC ;L or Kg ; blood flow to kidney (calculated)
QSI=QSIc*QC ;L or Kg ; blood flow to small intestine tissue (calculated)
QLI=QLIc*QC ;L or Kg ; blood flow to large intestine tissue (calculated)
;-------------------------------------------------------------------------------------------------------------------
; glomerular filtration rate
GFR = 125 ;mL/min ; GFR scaled to body weight (Levey et al.)
;=====================================================================
; Physicochemical parameters
;=====================================================================
; molecular weights (g/mol)
MWAME = 272.25 ; parent compound
; pKa1
pKa1AME =7.71 ; parent compound Chemaxon playground (v1.6.2)
; logP
logPAME = 3.32 ; parent compound Chemaxon playground (v1.6.2)
; logD (parent compound)
;logDapical = 3.32 Chemaxon playground (v1.6.2)
;logDbasal = 3.1 Chemaxon playground (v1.6.2)

; partition coefficients ;calculated by qivivetools.wu.nl (Berezhkovskiy algorithm)
; AME in main model
PLAME =3.8 ; liver/blood partition coefficient
PRAME = 0.982 ; rapidly perfused tissue/blood partition coefficient
PSAME = 5.509 ; slowly perfused tissue/blood partition coefficient
PFAME = 7.1 ; fat/blood partition coefficient
PMAME = 2.44 ; muscle/blood partition coefficient
PKAME = 2.41 ; kidney/blood partition coefficient
FUAME = 0.058 ; faction unbound plasma according to Lobell&Sivarajah
PSIAME =4.72 ; gut/blood partition coefficient
;-------------------------------------------------------------------------------------------------------------------
; absorption/transfer rates
logPappAME = -5.11 ;log(cm/s) ;log permeability estimated

```
PeffAME = (3600/10)*(10^(0.6836*LogPappAME- 0.5579))    ;dm/h    ;scaled according to Sun et al.


KaAME = PeffAME*areaSI    ;L/h ;transfer rate of AME from SI lumen to gut
KbAME = PeffAME*areaLI    ;L/h ;transfer rate of AAME from LI lumen to gut

; gastric emptying (kelst)
GEstc = 11.8    ; (min) half time of meal in mice stomach
GEst = GEstc /60    ; (h)
LNhalf= -0.69314718    ; LN(1/2)
kelST =  -LNhalf/ GEst    ; (h-1) constant of gastric excretion    ; from Ct=C0*e^(-kel*t)

tpassSI = 4    ;h    ;passing time through small intestine (ICRP)
Ksi = 1/tpassSI    ;/h    ;SI to LI transfer constant
tpassLI = 36    ;h    ;passing time through large intestine (ICRP)
Kli = 1/tpassLI    ;/h    ;LI to excretion transfer constant
;=================================================================
; Kinetic parameters
;=================================================================
;metabolism of rat liver
; scaling factors
VLS9 = 107.3    ;mg S9 protein/gram liver    ; liver S9 protein yield (Wang et al. 2019)
VLMP = 32    ;mg protein/gram liver    ; liver microsomal protein yield (Barter et al. 2007)

;Phase 1 metabolism
KmLAME = 29.4    ;µmol/L    ;Michaelis constant of hepatic phase I metabolism (Borsos et al)
VmaxLAMEc = 0.000772    ;µmol/mg/min   microsomal protein (Borsos et al)
VmaxLAME = VmaxLAMEc*VLMP*VL*60*10^3    ;µmol/h

;Phase 2 glucuronidation

KmLAMEgluc = 0.97    ;µmol/L    ;Michaelis constant of hepatic glucuronidation (in vitro)
VmaxLAMEglucc =0.0105    ;µmol/(mg*min) (in vitro)
VmaxLAMEgluc = VmaxLAMEglucc*VLS9*VL*60*10^3    ;µmol/h
;=================================================================
; Run settings
;=================================================================
;-----------------------------------------------------------------------------------------------------------
; oral dose
ODOSEAME = 0.0014*BW ;µg    ;oral dose alternariol, variable
AODOSEAME = ODOSEAME/MWAME    ;umol    ;oral dose AME in umol
;=================================================================
; Main model calculations/dynamics:
;=================================================================
;-----------------------------------------------------------------------------------------------------------
;ASIAME: amount of AME in small intestinal lumen compartment    ;µmol
CSIAME=ASIAME/VSIL    ;umol/L
;-----------------------------------------------------------------------------------------------------------
;large intestine compartment
;ALIAME: amount of AME in large intestine lumen    ;µmol
CLIAME = ALIAME/VLIL    ;umol/L
;-----------------------------------------------------------------------------------------------------------
;gut small intestine tissue compartment
;AGSIAME: amount of AME in small intestine tissue    ;µmol
CGSIAME = AGSIAME/VSI    ;µmol/L
CVGSIAME = CGSIAME/PSIAME
;-----------------------------------------------------------------------------------------------------------
; liver compartment
; ALAME: amount of AME in liver ;µmol
CLAME = ALAME/VL    ;umol/L
CVLAME = CLAME/PLAME    ;partition with liver tissue
;-----------------------------------------------------------------------------------------------------------
; portal vein perfused tissue compartment
; APAME = amount of AME in spleen tissue ;µmol
```

```
CPAME = APAME/VP                    ;umol/L
;-----------------------------------------------------------------------------------------------------------------
; fat (adipose tissue) compartment
CFAME = AFAME/VF                    ;umol/L
CVFAME = CFAME/PFAME                            ;partition with adipose tissue
;-----------------------------------------------------------------------------------------------------------------
; muscle compartment
; AMAME = amount of AME in muscle tissue    ;µmol
CMAME = AMAME/VM                    ;umol/L
CVMAME = CMAME/PMAME                            ;partition with muscle tissue
;-----------------------------------------------------------------------------------------------------------------
; rapidly perfused tissue
CRAME = ARAME/VR                    ;umol/L
CVRAME = CRAME/PRAME                            ;partition with muscle tissue
;----------------------------------------------------------------------------------------------------------------
; slowly perfused tissue
CSAME = ASAME/VS                    ;umol/L
CVSAME = CSAME/PSAME                            ;partition with slowly perfused tissue
;-----------------------------------------------------------------------------------------------------------------
; kidney
CKAME = AKAME/VK                    ;umol/L
CVKAME = CKAME/PKAME                            ;partition with kidney tissue
;-----------------------------------------------------------------------------------------------------------------
; blood compartment
CBAME=ABAME/VB                      ;umol/LCVBAME = CBAME*FUAME                     ;free concentration in
plasma
CBAMEug = CBAME*MWAME        ;µg/L = ng/mL                    ;concentration in blood in ng/ml
;-----------------------------------------------------------------------------------------------------------------
;=================================================================
; Main model: mass balance calculation
;================================================================
Total = AODOSEAME
Calculated = ASTAME + ASIAME  + ALIAME + AGSIAME  + ALAME+ ABAME+ AKAME + ARAME + ASAME +
AFAME + APAME + AMAME + AUCAMEUr +FECAME+exAMEmetab + exAMEglucur

ERROR=((Total-Calculated)/(Total+1E-30))*100
MASSBAL=Total-Calculated + 1

{End Globals}
```
