## Supplementary material for "Human internal exposures to alternariol and its monomethyl ether are predicted below thresholds of *in vitro* toxicity by physiologically based kinetic modeling": S8 Female model for AME.pdf

**Model S8:** Female model for AME

```
METHOD RK4

STARTTIME = 0
STOPTIME = 10
DT = 0.00002

{Type Equations Here.}
{Top model}
{Reservoirs}

d/dt (ASTAME) = - transSTtoSI + oraldose
INIT ASTAME = 0
LIMIT ASTAME >= 0
d/dt (ASIAME) = -transSItoSIG - transSItoLI  +  transSTtoSI
INIT ASIAME = 0
LIMIT ASIAME >= 0
d/dt (ALIAME) =  +  transSItoLI - fecalex - transLItoL
INIT ALIAME = 0
LIMIT ALIAME >= 0
d/dt (AGSIAME) =  + transSItoSIG  -venGS
INIT AGSIAME = 0
LIMIT AGSIAME >= 0
d/dt (FECAME) = + fecalex
INIT FECAME = 0
LIMIT FECAME >= 0
d/dt (ALAME) = - venL + artL + venP + venGS  + transLItoL - Phase1metab - AMEglucur
INIT ALAME = 0
LIMIT ALAME >= 0
d/dt (ABAME) = + venL + venS + venR + venF + venK + venM  - artL - artP - artS - artR -artF - artK - artM
INIT ABAME = 0
LIMIT ABAME >= 0
d/dt (APAME) = + artP - venP
INIT APAME = 0
LIMIT APAME >= 0
d/dt (ASAME) = + artS - venS
INIT ASAME = 0
LIMIT ASAME >= 0
d/dt (ARAME) = + artR - venR
INIT ARAME = 0
LIMIT ARAME >= 0
d/dt (AFAME) = + artF - venF
INIT AFAME = 0
LIMIT AFAME >= 0
d/dt (AUAME) = + artU - venU
INIT AUAME = 0
LIMIT AUAME >= 0
d/dt (AKAME) = + artK - venK - urineAME
INIT AKAME = 0
LIMIT AKAME >= 0
d/dt (AUCAMEUr) = + urineAME
INIT AUCAMEUr = 0
d/dt (AMAME) = + artM - venM
INIT AMAME = 0
LIMIT AMAME >= 0
d/dt (Start) = - oraldose
INIT Start = AODOSEAME
LIMIT Start >=0
d/dt (exAMEmetab) = + Phase1metab
INIT exAMEmetab = 0
LIMIT exAMEmetab >= 0
d/dt (exAMEglucur) = + AMEglucur
INIT exAMEglucur = 0
```

```
LIMIT exAMEglucur >= 0

{Flows}
 ; arterial (art) and venous (ven) blood flows :
venL = QL*CVLAME
artL = (QL-QP)*CBAME
venP = QP*CPAME
artP = QP*CBAME
venS = QS*CVSAME
artS = QS*CBAME
venR = QR*CVRAME
artR = QR*CBAME
venF = QF*CVFAME
artF = QF*CBAME
venM = QM*CVMAME
artM = QM*CBAME
venK = QK*CVKAME
artK = QK*CBAME
venGS = QSI*CVGSIAME
artGS = QSI*CBAME
venU=QU*CVUAME
artU=QU*CBAME

 ; excretion :
urineAME = GFR*CVKAME*60/1000  ;  (L/h)
fecalex = Kli*ALIAME

; gastrointestinal tract :
transSTtoSI = kelST * ASTAME                    ; from stomach lumen to small intestine lumen
transSItoSIG = KaAME*CSIAME                     ; from small intestine lumen to gut small intestine tissue
transSItoLI = ASIAME*Ksi                        ; from small intestine lumen to large intestine lumen
transLItoL = KbAME*CLIAME                       ; from large intestine lumen to liver tissue

oraldose = PULSE(AODOSEAME, 0, 24)

;Liver metabolism :
Phase1metab = VmaxLAME * CVLAME/(KmLAME + CVLAME)          ; MichaelisMenten
AMEglucur = VmaxLAMEgluc * CVLAME/(KmLAMEgluc + CVLAME)  ; MichaelisMenten

{Globals}

;AME female model
;=====================================================================
; Physiological parameters
;=====================================================================
;bodyweight
BW = 60 ;Kg          ; avg. BW in validation study (ICRP)
;-------------------------------------------------------------------------------------------------------------------
; relative tissue volumes
; fraction of BW
VLc = 0.0298                    ; fraction of liver tissue (ICRP)
VPc = 0.0064                    ; fraction portal vein perfused tissue (stomach, spleen, pancreas) tissue (ICRP)
VRc =0.0199                     ; fraction of rapidly perfused tis. (Q/V >>1, heart + lungs) tissue (ICRP)
VSc = 0.1748                    ; fraction of slowly perfused tissue (bone, skin) tissue (ICRP)
VFc = 0.3055                    ; fraction of fat tissue (adipose) tissue (ICRP)
VMc = 0.2985                    ; fraction of muscle tissue tissue (ICRP)
VKc = 0.0059                    ; fraction of kidneys tissue (ICRP)
VSIc = 0.0125                   ; fraction of small intestine tissue (ICRP)
VLIc = 0.0074                   ; fraction of large intestine tissue (ICRP)
VBc = 0.065                     ; fraction of blood (ICRP)
VUc= 0.0016                     ;fraction of uterus tissue (ICRP)

;-------------------------------------------------------------------------------------------------------------------
; calculated tissue volumes
```

```
VL = VLc*BW              ;L or Kg                ; volume of liver tissue (calculated)
VP = VPc*BW              ;L or kg                ; volume of portal vein perfused tissue
VR = VRc*BW              ;L or Kg                ; volume of rapidly perfused tissue (calculated)
VS = VSc*BW              ;L or Kg                ; volume of slowly perfused tissue (calculated)
VF = VFc*BW              ;L or Kg                ; volume of fat tissue (calculated)
VM = VMc*BW              ;L or Kg                ; volume of muscle tissue (calculated)
VK = VKc*BW              ;L or Kg                ; volume of kidney (calculated)
VB = VBc*BW              ;L or Kg                ; volume of blood (Brown et al, calculated)
VSI = VSIc*BW            ;L or Kg                ; volume of small intestine tissue (calculated)
VLI = VLIc*BW            ;L or Kg                ; volume of large intestine tissue  (calculated)
VU = VUc*BW              ;L or Kg                ; volume of uterus tissue (calculated)
VSIL = 4.13              ;L or Kg                ; volume of small intestine lumen (ICPR, calculated)
VLIL = 2.16              ;L or Kg                ; volume of large intestine lumen (ICPR, calculated)
;-----------------------------------------------------------------------------------------------------------------
;surface areas
areaSI = 2709.066        ;dm2          ;surface area of SI lumen (Helander et al.)
areaLI = 98.018          ;dm2          ;surface area of LI lumen (Helander et al.)
;-----------------------------------------------------------------------------------------------------------------
;Blood flow rates,
QCc = 5.9                ; (L/min), cardiac output (ICRP)
QPc = 0.05               ; fraction of blood flow to P (spleen, stom., pancreas) (ICRP)
QLc = 0.065              ; fraction of blood flow to liver in total (ICRP)
QRc = 1.05               ; fraction of blood flow to rapidly perfused tissue (heart+lungs)     (ICRP)
QSc = 0.1                ; fraction of blood flow to slowly perfused tissue (skin+Bones)       (ICRP)
QFc = 0.085              ; fraction of blood flow to fat  (ICRP)
QMc =0.12                ; fraction of blood flow to muscle (ICRP)
QKc =0.17                ; fraction of blood flow to kidneys (ICRP)
QSIc =0.11               ; fraction of blood flow to small intestine (ICRP)
QLIc=0.05                ; fraction of blood flow to large intestine (ICRP)
QUc= 0.004               ;fraction of blood to uterus (ICRP)

QC = QCc*60              ; (L/h)       ; cardiac output (calculated)
QP = QPc*QC              ;L or Kg      ; blood flow to portal vein perfused tissue (calculated)
QL = QLc*QC              ;L or Kg      ; blood flow to liver total (calculated)
QR = QRc*QC              ;L or Kg      ; blood flow to rapidly perfused tissue (calculated)
QS = QSc*QC              ;L or Kg      ; blood flow to slowly perfused tissue (calculated)
QF = QFc*QC              ;L or Kg      ; blood flow to fat tissue (calculated)
QM = QMc*QC              ;L or Kg      ; blood flow to muscle tissue (calculated)
QK = QKc*QC              ;L or Kg      ; blood flow to kidney (calculated)
QSI=QSIc*QC              ;L or Kg      ; blood flow to small intestine tissue (calculated)
QLI=QLIc*QC              ;L or Kg      ; blood flow to large intestine tissue (calculated)
QU = QUc*QC              ;L or Kg      ; blood flow to uterus tissue (calculated)
;-----------------------------------------------------------------------------------------------------------------
; glomerular filtration rate
GFR = 125       ;mL/min       ; GFR scaled to body weight (Levey et al.)
;================================================================
; Physicochemical parameters
;================================================================
; molecular weights (g/mol)
MWAME = 272.25           ; parent compound
; pKa1
pKa1AME =7.71 ; parent compound Chemaxon playground (v1.6.2)
; logP
logPAME = 3.32 ; parent compound Chemaxon playground (v1.6.2)
; logD (parent compound)
;logDapical = 3.32 Chemaxon playground (v1.6.2)
;logDbasal = 3.1 Chemaxon playground (v1.6.2)

; partition coefficients            ;calculated by qivivetools.wu.nl (Berezhkovskiy algorithm)
; AME in main model
PLAME =3.8               ; liver/blood partition coefficient
PRAME = 0.982            ; rapidly perfused tissue/blood partition coefficient
PSAME = 5.551            ; slowly perfused tissue/blood partition coefficient
PFAME = 7.1              ; fat/blood partition coefficient
```

```
; liver compartment
; ALAME: amount of AME in liver ;µmol
CLAME = ALAME/VL                ;umol/L
CVLAME = CLAME/PLAME                    ;partition with liver tissue
;--------------------------------------------------------------------------------------------------------------------
; portal vein perfused tissue compartment
; APAME = amount of AME in spleen tissue ;µmol
CPAME = APAME/VP                ;umol/L
;--------------------------------------------------------------------------------------------------------------------
; fat (adipose tissue) compartment
CFAME = AFAME/VF                ;umol/L
CVFAME = CFAME/PFAME                    ;partition with adipose tissue
;--------------------------------------------------------------------------------------------------------------------
; muscle compartment
; AMAME = amount of AME in muscle tissue    ;µmol
CMAME = AMAME/VM                ;umol/L
CVMAME = CMAME/PMAME                    ;partition with muscle tissue
;--------------------------------------------------------------------------------------------------------------------
; rapidly perfused tissue
CRAME = ARAME/VR                ;umol/L
CVRAME = CRAME/PRAME                    ;partition with muscle tissue
;-------------------------------------------------------------------------------------------------------------------
; slowly perfused tissue
CSAME = ASAME/VS                ;umol/L
CVSAME = CSAME/PSAME                    ;partition with slowly perfused tissue
;--------------------------------------------------------------------------------------------------------------------
; kidney
CKAME = AKAME/VK                ;umol/L
CVKAME = CKAME/PKAME                    ;partition with kidney tissue
;--------------------------------------------------------------------------------------------------------------------
; blood compartment
CBAME=ABAME/VB                  ;umol/LCVBAME = CBAME*FUAME                ;free    concentration    in
plasma
CBAMEug = CBAME*MWAME       ;µg/L = ng/mL               ;concentration in blood in ng/ml
;--------------------------------------------------------------------------------------------------------------------
; uterus compartment
CUAME=AUAME/VU                  ;umol/L
CVUAME = CUAME*PMAME                    ;free concentration in plasma
;------------------------------------------------------------------------------------------------------------------
;=====================================================================
; Main model: mass balance calculation
;====================================================================
Total = AODOSEAME
Calculated = ASTAME + ASIAME  + ALIAME + AGSIAME  + ALAME+ ABAME+ AKAME + ARAME + ASAME +
AFAME + APAME + AMAME + AUCAMEUr +FECAME+exAMEmetab + exAMEglucur + AUAME

ERROR=((Total-Calculated)/(Total+1E-30))*100
MASSBAL=Total-Calculated + 1

{End Globals}
```
