## Supplementary material for "Human internal exposures to alternariol and its monomethyl ether are predicted below thresholds of *in vitro* toxicity by physiologically based kinetic modeling": S10 Male toddler model for AME.pdf

```
VB = VBc*BW              ;L or Kg              ; volume of blood (Brown et al, calculated)
VSI = VSIc*BW            ;L or Kg              ; volume of small intestine tissue (calculated)
VLI = VLIc*BW            ;L or Kg              ; volume of large intestine tissue  (calculated)
VSIL = 0.34              ;L or Kg              ; volume of small intestine lumen (ICPR, calculated)
VLIL = 0.228             ;L or Kg              ; volume of large intestine lumen (ICPR, calculated)
;-------------------------------------------------------------------------------------------------------------
;surface areas
areaSI = 1250.338        ;dm2          ;surface area of SI lumen (Helander et al.)
areaLI = 58.81           ;dm2          ;surface area of LI lumen (Helander et al.)
;-------------------------------------------------------------------------------------------------------------
;Blood flow rates,
QCc = 1.2          ; (L/min), cardiac output (ICRP)
QPc = 0.05         ; fraction of blood flow to P (spleen, stom., pancreas) (ICRP)
QLc = 0.065        ; fraction of blood flow to liver in total (ICRP)
QRc = 1.04         ; fraction of blood flow to rapidly perfused tissue (heart+lungs)      (ICRP)
QSc = 0.1          ; fraction of blood flow to slowly perfused tissue (skin+Bones)        (ICRP)
QFc = 0.05         ; fraction of blood flow to fat  (ICRP)
QMc =0.17          ; fraction of blood flow to muscle (ICRP)
QKc =0.19          ; fraction of blood flow to kidneys (ICRP)
QSIc =0.1          ; fraction of blood flow to small intestine (ICRP)
QLIc=0.04          ; fraction of blood flow to large intestine (ICRP)

QC = QCc*60        ; (L/h)              ; cardiac output (calculated)
QP = QPc*QC        ;L or Kg  ; blood flow to portal vein perfused tissue (calculated)
QL = QLc*QC        ;L or Kg  ; blood flow to liver total (calculated)
QR = QRc*QC        ;L or Kg  ; blood flow to rapidly perfused tissue (calculated)
QS = QSc*QC        ;L or Kg  ; blood flow to slowly perfused tissue (calculated)
QF = QFc*QC        ;L or Kg  ; blood flow to fat tissue (calculated)
QM = QMc*QC        ;L or Kg  ; blood flow to muscle tissue (calculated)
QK = QKc*QC        ;L or Kg  ; blood flow to kidney (calculated)
QSI=QSIc*QC        ;L or Kg  ; blood flow to small intestine tissue (calculated)
QLI=QLIc*QC        ;L or Kg  ; blood flow to large intestine tissue (calculated)
;-------------------------------------------------------------------------------------------------------------
; glomerular filtration rate
GFR = 125          ;mL/min              ; GFR scaled to body weight (Levey et al.)
;=============================================================================================================
; Physicochemical parameters
;=============================================================================================================
; molecular weights (g/mol)
MWAME = 272.25                ; parent compound
; pKa1
pKa1AME =7.71 ; parent compound Chemaxon playground (v1.6.2)
; logP
logPAME = 3.32 ; parent compound Chemaxon playground (v1.6.2)
; logD (parent compound)
;logDapical = 3.32 Chemaxon playground (v1.6.2)
;logDbasal = 3.1 Chemaxon playground (v1.6.2)

; partition coefficients                 ;calculated by qivivetools.wu.nl (Berezhkovskiy algorithm)
; AME in main model
PLAME =3.8               ; liver/blood partition coefficient
PRAME = 1.014            ; rapidly perfused tissue/blood partition coefficient
PSAME = 5.540            ; slowly perfused tissue/blood partition coefficient
PFAME = 7.1              ; fat/blood partition coefficient
PMAME = 2.44             ; muscle/blood partition coefficient
PKAME = 2.41             ; kidney/blood partition coefficient
FUAME = 0.058            ; faction unbound plasma according to Lobell&Sivarajah
PSIAME =4.72             ; gut/blood partition coefficient
;-------------------------------------------------------------------------------------------------------------
; absorption/transfer rates
logPappAME = -5.11                ;log(cm/s)                      ;log permeability estimated
PeffAME = (3600/10)*(10^(0.6836*LogPappAME- 0.5579))       ;dm/h    ;scaled according to Sun et al.
```

```
KaAME = PeffAME*areaSI              ;L/h ;transfer rate of AME from SI lumen to gut
KbAME = PeffAME*areaLI              ;L/h ;transfer rate of AAME from LI lumen to gut

; gastric emptying (kelst)
GEstc = 11.8                        ; (min) half time of meal in mice stomach
GEst = GEstc /60          ; (h)
LNhalf= -0.69314718                 ; LN(1/2)
kelST =  -LNhalf/ GEst    ; (h-1) constant of gastric excretion          ; from Ct=C0*e^(-kel*t)

tpassSI = 4       ;h      ;passing time through small intestine (ICRP)
Ksi = 1/tpassSI   ;/h     ;SI to LI transfer constant
tpassLI = 32              ;h       ;passing time through large intestine (ICRP)
Kli = 1/tpassLI   ;/h     ;LI to excretion transfer constant
;==========================================================================
; Kinetic parameters
;==========================================================================
;metabolism of rat liver
; scaling factors
VLS9 = 107.3      ;mg S9 protein/gram liver        ; liver S9 protein yield (Wang et al. 2019)
VLMP = 32         ;mg protein/gram liver           ; liver microsomal protein yield (Barter et al. 2007)

;Phase 1 metabolism
KmLAME = 29.4                       ;µmol/L          ;Michaelis constant of hepatic phase I metabolism (Borsos et al)
VmaxLAMEc = 0.000772                      ;µmol/mg/min   microsomal protein (Borsos et al)
VmaxLAME = VmaxLAMEc*VLMP*VL*60*10^3       ;µmol/h

;Phase 2 glucuronidation

KmLAMEgluc = 0.97                         ;µmol/L          ;Michaelis constant of hepatic glucuronidation (in vitro)
VmaxLAMEglucc =0.0105               ;µmol/(mg*min) (in vitro)
VmaxLAMEgluc = VmaxLAMEglucc*VLS9*VL*60*10^3        ;µmol/h
;==========================================================================
; Run settings
;==========================================================================
;--------------------------------------------------------------------------
; oral dose
ODOSEAME = 0.0579*BW ;µg                   ;oral dose AME, variable
AODOSEAME = ODOSEAME/MWAME          ;umol            ;oral dose AME in umol
;==========================================================================
; Main model calculations/dynamics:
;==========================================================================
;--------------------------------------------------------------------------
;ASIAME: amount of AME in small intestinal lumen compartment          ;µmol
CSIAME=ASIAME/VSIL                                            ;umol/L
;--------------------------------------------------------------------------
;large intestine compartment
;ALIAME: amount of AME in large intestine lumen                       ;µmol
CLIAME = ALIAME/VLIL                                          ;umol/L
;--------------------------------------------------------------------------
;gut small intestine tissue compartment
;AGSIAME: amount of AME in small intestine tissue                     ;µmol
CGSIAME = AGSIAME/VSI                                                 ;µmol/L
CVGSIAME = CGSIAME/PSIAME
;--------------------------------------------------------------------------
; liver compartment
; ALAME: amount of AME in liver ;µmol
CLAME = ALAME/VL                    ;umol/L
CVLAME = CLAME/PLAME                      ;partition with liver tissue
;--------------------------------------------------------------------------
; portal vein perfused tissue compartment
; APAME = amount of AME in spleen tissue ;µmol
CPAME = APAME/VP                    ;umol/L
;--------------------------------------------------------------------------
; fat (adipose tissue) compartment
```
